# Uridine cytidine kinases dictate the therapeutic response of molnupiravir via its bioactivation

**DOI:** 10.1101/2025.05.13.653844

**Authors:** Huazhang Shu, Sushma Sharm, Seher Alam, Lilian Frank, Marianna Tampere, André B.P. van Kuilenburg, Nicholas C.K. Valerie, Mikael Altun, Andrei Chabes, Sean G. Rudd, Si Min Zhang

## Abstract

Molnupiravir is a nucleoside analogue antiviral drug with activities against a broad spectrum of RNA viruses, including its clinical indication SARS-CoV-2. Whilst its antiviral mechanism-of-action is well defined, host pharmacogenetic factors that regulate its therapeutic responses – efficacy as well as selectivity – have not been mechanistically deciphered and characterized. Here we identified that uridine cytidine kinase (UCK), the first and rate-limiting kinase of the pyrimidine salvage pathway, could effectively phosphorylate N4-hydroxycytidine (NHC), the active compound of molnupiravir, and thereby dictate its anti-SARS-CoV-2 efficacy, as well as its selectivity index. Using target engagement and enzyme kinetic assays, we demonstrate that recombinant UCK isoform 1 (UCK1) and 2 (UCK2) effectively bind and phosphorylate NHC, with UCK2 displaying 9-fold higher catalytic efficiency. Accordingly, in SARS-CoV-2-infected cells, downregulation of UCK2 via siRNA hampered the intracellular accumulation of the triphosphate antiviral metabolite of NHC, resulting in 10-fold reduction of antiviral efficacy. Furthermore, we could recapitulate this using a pan-UCK small molecule inhibitor. Critically, both UCK downregulation and inhibition significantly reduced the selectivity index of molnupiravir/NHC. Altogether, this work underscores the pivotal roles of UCK enzymes in upholding molnupiravir efficacy and therapeutic window, and furthermore, highlights their potential as pharmacologically tractable targets for tailoring the therapeutic response to this antiviral agent.

## INTRODUCTION

Uridine and cytidine kinase (UCK) (EC 2.7.1.48) is a pyrimidine ribonucleoside kinase with two isoforms sharing ∼70% sequential homology. Isoform 1 (UCK1) is uniformly expressed across tissues, whilst the catalytically superior isoform 2 (UCK2) is primarily reserved for embryonic and placental tissues, but can be re-expressed in malignancies [1, 2]. Physiologically, UCKs governs the first and rate-limiting step of the pyrimidine salvage pathway by catalyzing the mono-phosphorylation of plasma-derived uridine and cytidine into uridine and cytidine monophosphates (UMP and CMP), respectively. UMP and CMP can then be sequentially phosphorylated by CMP kinase (CMPK) and nucleoside-diphosphate kinase (NDPK) into uridine triphosphate (UTP) and cytidine triphosphate (CTP), fueling cellular demand for pyrimidine nucleotides (**Fig. 1a**) [3].

**Figure 1.**
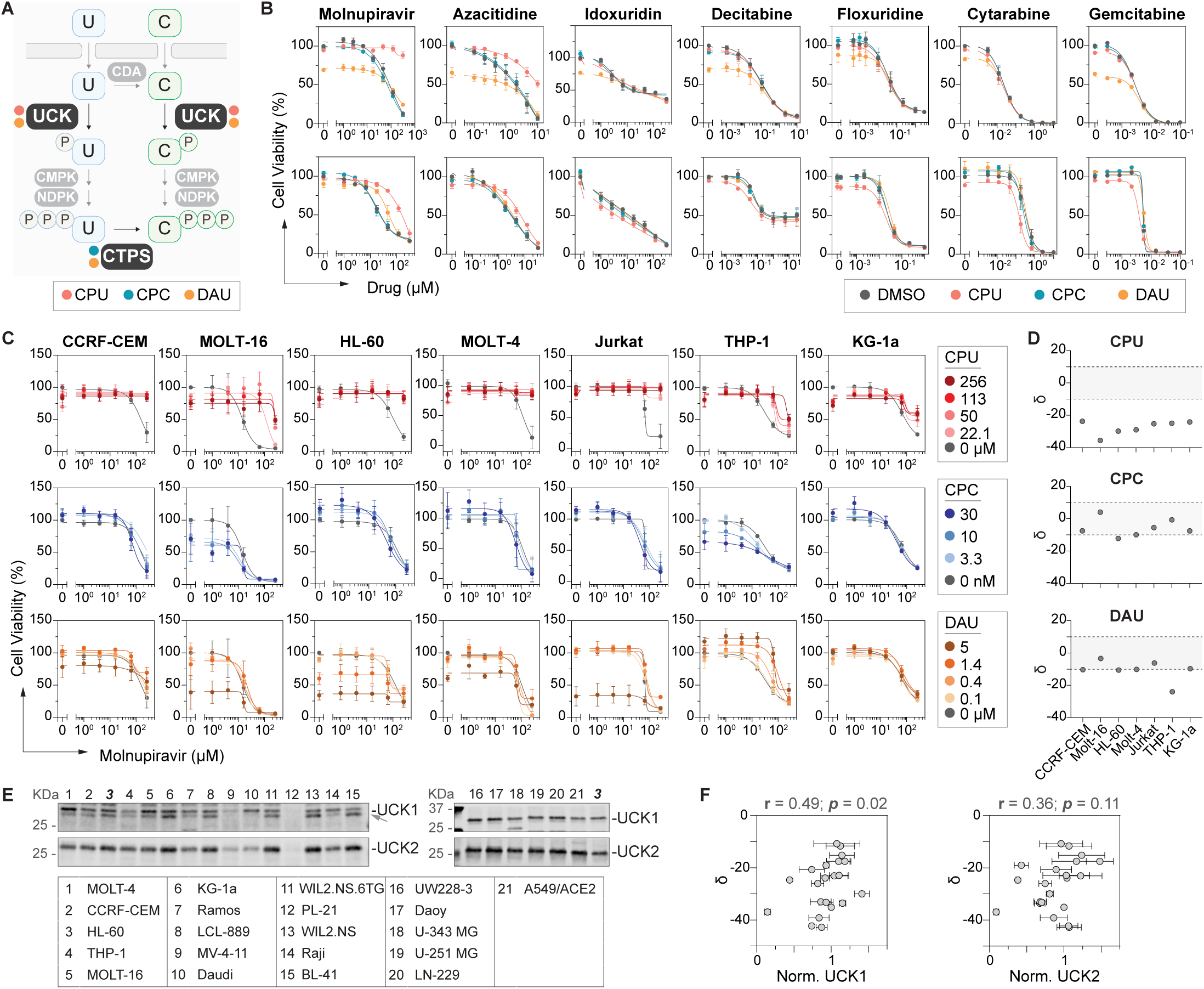
UCK inhibition limits cellular response to molnupiravir and its cell-active metabolite NHC. **A.** Schematic representation of cellular roles of UCK in pyrimidine salvage pathway. UCKs governs the first- and rate-limiting step of pyrimidine salvage pathway. It catalyzes the mono-phosphorylation of uridine and cytidine into uridine monophosphate (UMP) and cytidine monophosphate (CMP), respectively, which upon subsequent phosphorylation by CMPK and NDPK are converted to the tri-phosphorylated nucleotides. Cytidine triphosphate (CTP) could be further converted from uridine triphosphate (UTP) by CTPS. In the cell viability-based screening, UCK inhibitor CPU, UCK/CTPS dual inhibitor DAU, and finally CPTS inhibitor CPC were employed, as highlighted in the schematic representation. UCK, uridine cytidine kinase 1/2; CTPS, CTP synthase 1/2; CMPK, cytidine monophosphate kinase; CDA, cytidine deaminase; NDPK, nucleoside-diphosphate kinase; CPU, cyclopentenyl uracil; DAU, 3-Deazauridine; CPC, cyclopentenyl cytosine. **B.** Focused nucleoside analogue drug library screening identified UCK as a limiting factor for the cellular response of molnupiravir. AML HL-60 and THP1 cells were treated with nucleoside analogues drugs in the presence or absence of 100 µM CPU, 12 nM CPC, and 1 µM DAU for 96 hours before cell viabilities were determined using a resazurin reduction assay. Relative viability percentages were determined by normalizing background-subtracted fluorescence signals at 584nm to DMSO controls. In both HL-60 and THP1 cells, inhibition of UCK by CPU and DAU, but not CTPS inhibition by CPC, retarded the cytotoxicity of molnupiravir and azacitidine. **C-D.** Inhibition of UCK consistently antagonized NHC, metabolite of molnupiravir, in multiple cell lines. In C, relative cell viability curves in multiple leukemia cell lines treated with NHC, in the presence or absence of CPU, CPC, or DAU; in D, the relative cell viability data as presented in C were further used to determine synergy scores (δ), calculated using SynergyFinder. **E-F.** Cellular UCK levels positively correlated with CPU-induced NHC antagonism. In E, Western blot analysis of UCK expression levels in multiple cell lines, where lysates containing equal amount of protein were separated by SDS-PAGE, probed with UCK1 and UCK2 specific antibodies, and then subject to densitometry analysis. UCK1 and UCK2 expression in the cell lines were normalized to the levels in HL-60 cells, and were presented as relative levels. In F, cell lines in E were subject to resazurin reduction assay upon 96-hour treatment with NHC or DMSO control, in the presence or absence of CPU. Synergy scores were subsequently determined based on the cell viability data, as described in D, and further demonstrated positive correlations with both UCK1 and UCK2 expression levels.

Nucleotide salvage pathways are also central for the activation of nucleoside analogue (NA) therapeutics, which constitute an important class of anticancer and antiviral agents. NA drugs require stepwise phosphorylation to generate their active metabolites, typically their triphosphate (NA-TP) forms, which can then compete with their physiological counterparts to exert their therapeutic effects. Similar to its physiological activity, UCKs have been shown to mono-phosphorylate several uridine and cytidine analogues including azacitidine and 5-fluorouridine and the investigational agent RX-3117, and thereby control the accumulation of the active NA-TP pool [1, 4, 5]. Furthermore, low UCK expression and loss-of-activity mutations have been associated with resistance to azacitidine, which can be restored upon UCK re-expression [6–9]. Whilst these studies support UCKs as putative biomarkers for azacitidine therapy, the broader role of UCKs in governing the efficacy of NA-based therapies, and particularly newly approved NA drugs, such as the severe acute respiratory coronavirus 2 (SARS-CoV-2) therapy molnupiravir [10], remains unclear.

Molnupiravir is the isopropyl ester pro-drug of β-D-N4-hydroxycytidine (NHC), a broad-spectrum antiviral with activities against multiple RNA viruses aside from its clinical indication SARS-CoV-2, including chikungunya virus [11], hepatitis C virus [12], influenza viruses [13], Ebola virus [14], and other coronavirus family members including SARS-CoV-1 [15]. Upon administration, molnupiravir is rapidly converted to NHC by plasma esterase, which is then taken up into cells and subsequently phosphorylated to NHC triphosphate (NHC-TP). As a close analogue of CTP, NHC-TP is readily mis-incorporated into the viral genome by the viral RNA polymerase, and is then ambiguously matched with G or A in subsequent rounds of replication, resulting in defective virions with lethal mutation burden and abolished infectivity [16]. Aside from introducing mutations in the viral genome, several studies have also shown that molnupiravir/NHC could induce mutations in the host cell genome in a similar fashion [17, 18]. A study by Xu et al utilizing genome-wide CRISPR/Cas9 screening has identified that UCK, particularly UCK2, is critical to the host mutagenesis activity of molnupiravir/NHC [18]. However, no direct and mechanistic characterization of UCKs, either UCK1 or UCK2, in the bioactivation of NHC and subsequent modulation of the antiviral therapeutic efficacy/selectivity have been reported.

In light of the translational potential of UCKs as biomarkers for NA therapy, we set out to systematically profile NA drugs phosphorylated and thereby activated by UCKs using a cell viability-based screening platform. We then identified NHC/molnupiravir as a potential substrate of UCKs. Subsequent target engagement studies, enzymatic activity assays, and intracellular drug metabolite measurements elucidated that both UCK1 and 2 could effectively phosphorylate NHC, with UCK2 being the preferred kinase and able to dictate the intracellular accumulation of NHC-TP. Accordingly, both UCK knockdown via siRNA and inhibition via cyclopentenyl uracil (CPU) effectively limited the anti-SARS-CoV-2 efficacy of NHC by 10 fold, which, unexpectedly, is further accompanied by a narrowed selectivity index. Collectively, these data establish the pivotal role of UCKs as the first step of NHC bioactivation and thereby the abilities of these enzymes to control drug efficacy and therapeutic window, and furthermore, underscore their potential as a viable target for tailoring NA drug response.

## RESULTS

### UCKs dictate molnupiravir response in a cell viability-based screening platform

Previous independent studies have shown that UCKs can bioactivate several NA oncology drugs. Here we initiated the study by systematically profiling NA drugs phosphorylated and thereby activated by UCKs, focusing on newly approved and emerging therapeutics with indications for oncology and/or viral infection. Firstly, a structure-based *in silico* screening of 3323 compounds was performed, resulting in a focused group of 27 compounds as potential substrate of UCKs. Albeit being optimized for selectivity index, antiviral NA drugs could cause cytotoxicity at high concentrations, which often directly correlates to the cellular levels of their triphosphate active metabolites [19]. Based on this observation, for the in vitro validation of our *in silico* screening output – either anti-viral or -cancer applications – we conducted a cell viability-based screening platform, where involvement of UCK in drug activation was assessed based on drug-induced cytotoxicity (**Fig. 1a**). Specifically, acute myeloid leukemia cell lines HL-60 and THP-1 were treated with increasing concentrations of screening drugs for 96 hours before cell viability was determined by resazurin reduction assay, done in the presence or absence of high doses of previously reported cell-active inhibitors of UCKs - cyclopentenyl uracil (CPU) [20] or 3-deazauridine (DAU) [21, 22]. Aside from UCKs, DAU primarily targets CTP synthase upon intracellular tri-phosphorylation. To specifically assess the influence of UCKs on drug efficacy, the screening campaign hence further included cyclopentenyl cytidine (CPC), a sub-micromolar CTP synthase inhibitor [23], as a control compound (**Fig. 1a**).

Using this assay, we observed that UCK inhibitors CPU and DAU, but not the CTP synthase inhibitor CPC, potently protected both HL-60 and THP-1 cells from azacitidine, an anti-leukemic agent known to be activated and thereby controlled by UCKs [6, 7], hence validating the screening setup. Interestingly, similar rescue was observed for the anti-SARS-CoV-2 drug molnupiravir (**Fig. 1b**). Molnupiravir is the isopropyl ester pro-drug of the phosphorylatable antiviral NHC, which is rapidly converted by plasma esterase upon administration of molnupiravir [24]. Confirming the screening result using molnupiravir, dose-response matrixes of NHC and DAU/CPU/CPC in multiple cell line models revealed that inhibition of UCKs by CPU consistently antagonized NHC-induced cytotoxicity with drug combination scores routinely below -20 (i.e. strong antagonism), which is followed by DAU but not CPC (**Fig. 1c-d**). These data suggest that UCKs are the principal kinases governing the intracellular responses of NHC/molnupiravir.

UCKs have two isoforms, the ubiquitously expressed UCK1 of lower catalytic efficacy and the more efficient UCK2 primarily reserved for placenta and overexpressed in cancers [1]. As the first step to delineate the principal UCK governing the cellular response of molnupiravir, we then correlated the intracellular UCK1 and UCK2 levels with CPU-induced NHC antagonism, in a panel of 21 cell lines of different tissue and disease origins (**Fig. 1e-f, Fig. S1**). We reason that for the enzyme critical for NHC bioactivation, upon its inhibition, cells of higher enzyme level would sustain greater loss of active drug metabolites and thereafter cytotoxicity. Indeed, we observed that in the tested cell lines, CPU-induced antagonism of NHC positively correlated with UCK1 protein levels (r=0.49; p=0.02). A similar positive correlation was observed with UCK2, albeit not significant. These data collectively suggest that UCK1 and UCK2 play principal roles in molnupiravir/NHC bioactivation and efficacy.

### NHC engages recombinant UCK1 and 2 in an ATP-dependent manner

To delineate mechanistically the roles of UCK1 and UCK2 in NHC metabolism, we first sought to confirm direct interactions between NHC and recombinant UCK1 and UCK2 using differential scanning fluorimetry (DSF). In the assay, ligand binding to recombinant proteins is interrogated based on the change of the protein thermal stability, as reflected by the shift of protein melting temperature (Tm). Despite sequence homology, recombinant UCK1 and UCK2 displayed drastically different Tms of approximately 60°C and 50°C, respectively. Nevertheless, both UCK1 and UCK2 were significantly stabilized in the presence of ATP by ∼10 °C, which further served as a prerequisite to their interactions, exemplified by further destabilization with known substrates – cytidine and azacitidine (**Fig. 2a-d**). Meanwhile, no such engagement was observed with the non-substrate thymidine (**Fig. S2a-b**), supporting the application of DSF to identify UCK interaction with substrates and cofactors.

**Figure 2.**
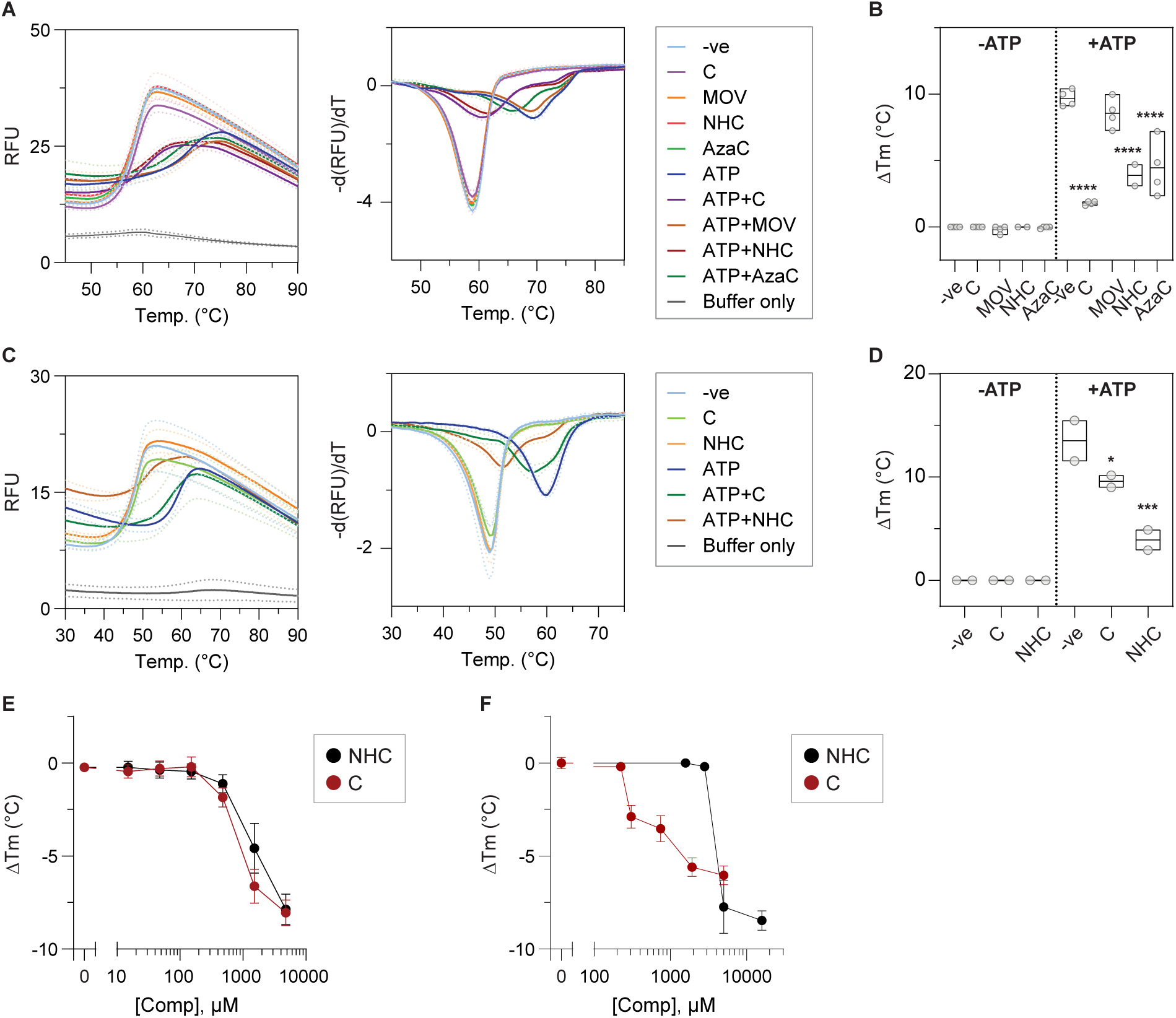
UCK1 and UCK2 preferentially interact with molnupiravir metabolite NHC in their ATP-bound state, similar to other known UCK substrates. **A-B.** Melting profiles of recombinant UCK1 in the presence of potential ligands, determined using differential scanning fluorometry (DSF) assays. Recombinant UCK1 (5 µM) was mixed with 5 mM molnupiravir (MOV), its phosphorylatable metabolite NHC, and known substrates azacitidine (AzaC) or cytidine (C), in the presence or absence of 5 mM ATP. Subsequently, protein thermal stability was examined using DSF. Mean fluorescence signals (solid line) ± SEM (dashed line) of a representative experiment performed in quadruplicate are shown in (A, *left panel*); melting temperatures (Tm) were then determined as the minima of negative derivative of the melting curve (A, *right panel*). In B, mean change of Tm (ΔTm) compared to protein only group of n=4 independent experiments are shown. Similar to the known substrates of UCK1 cytidine and azacitidine, the phosphorylatable metabolite of molnupiravir NHC, but not molnupiravir, effectively engaged recombinant UCK1 upon the addition of ATP, evidence by significant Tm changes compared to UCK1 bound to ATP only. Ordinary one-way ANOVA tests (Bonferroni’s multiple comparisons tests) were performed across treatment groups [ΔTm (protein only+ATP) vs. ΔTm (protein only+C), p < 0.0001, t = 12.73, DF = 26; ΔTm (protein only+ATP) vs. ΔTm (protein only+MOV), p = 0.3244, t = 1.815, DF = 26; ΔTm (protein only+ATP) vs. ΔTm (protein only+NHC), p < 0.0001, t = 7.641, DF = 26; ΔTm (protein only+ATP) vs. ΔTm (protein only+AzaC), p < 0.0001, t = 8.469, DF = 26], where asterisks signify statistical significance (∗∗∗∗p ≤ 0.0001). **C-D.** Melting profiles of recombinant UCK2 in the presence of potential ligands, determined using DSF assays as described in A-B. Recombinant UCK2 (3.5 µM) was incubated with 5mM compounds alone or in the presence of 5mM ATP, before protein thermal stabilities were determined using DSF. Similar to UCK1, NHC effectively engaged recombinant UCK2, but only in the presence of ATP. Mean fluorescence signals (solid line) ± SEM (dashed line) of a representative experiment performed in quadruplicate are shown (C, *left panel*); melting temperatures (Tm) were then determined as the minima of negative derivative of the melting curve (C, *right panel*). In D, mean change of Tm (ΔTm) compared to protein only group of n=2 independent experiments performed in triplicate are shown. Ordinary one-way ANOVA tests (Bonferroni’s multiple comparisons tests) were performed across treatment groups [ΔTm (protein only+ATP) vs. ΔTm (protein only+C), p = 0.048, t = 3.00, DF = 6; ΔTm (protein only+ATP) vs. ΔTm (protein only+NHC), p = 0.0007, t = 7.343, DF = 6], where asterisks signify statistical significance (∗p ≤ 0.05, ∗∗p ≤ 0.01, ∗∗∗p ≤ 0.001, ∗∗∗∗p ≤ 0.0001). **E-F.** Change of Tm of ATP-bound UCK1 (E) and UCK2 (F), in the presence of increasing concentrations of NHC or C, determined using DSF. Recombinant UCK1 and UCK2 were incubated with 5mM ATP, alone or together with increasing concentrations of NHC or C, before protein Tm were determined using DSF as described in the previous sections. Change of Tm (ΔTm) was calculated by normalizing to protein + ATP only group. Mean ΔTm ± SEM of n=4 independent experiments performed in quadruplicate are shown.

Both recombinant UCK1 and UCK2 were able to engage NHC, but only in the presence of ATP, similarly as the substrates cytidine and azacitidine (**Fig. 2a-d**). This was subsequently confirmed under dose-response setup, validating direct interaction between NHC and ATP-bound UCK1/2. Noticeably, with UCK1, NHC demonstrated comparable Tm shifts, suggesting similar binding affinity, as the canonical substrate cytidine, but drastically inferior to cytidine in UCK2 (**Fig. 2e-f, Fig. S2c-f**). Collectively, these data support that UCK1 and UCK2 dictated NHC cellular response through direct phosphorylation and activation.

### NHC is a substrate of UCK1 and UCK2

We next directly interrogated NHC as a substrate of UCK1 and UCK2, using an ADP-Glo-coupled kinase activity assay [25]. In the assay, the kinase reaction was coupled to the ADP-Glo™ detection system (Promega) that converts reaction-generated ADP into measurable luciferase signal, thereby allowing study of reaction kinetics. Using cytidine as the substrate, the ADP-Glo-coupled kinase activity assay produced kinetic parameters comparable to previous studies for both UCK1 and UCK2 [1], with UCK2 having superior catalytic efficiency of 56666.7 s^-1^·M^-1^ (**Fig. 3a-b, Fig. S3a-d**). We were further able to show that NHC was a substrate for both UCK1 and UCK2, with kcat of 53.4 ± 6.1 min^-1^ and 98.3 ± 9.3 min^-1^, and Km of 1.14 ± 0.32mM and 0.26 ± 0.07 mM, thereby specificity constant of 780.7 and 6301.3 s^-1^·M^-1^, respectively (**Fig. 3c-d, Fig. S3e-f**).

**Figure 3.**
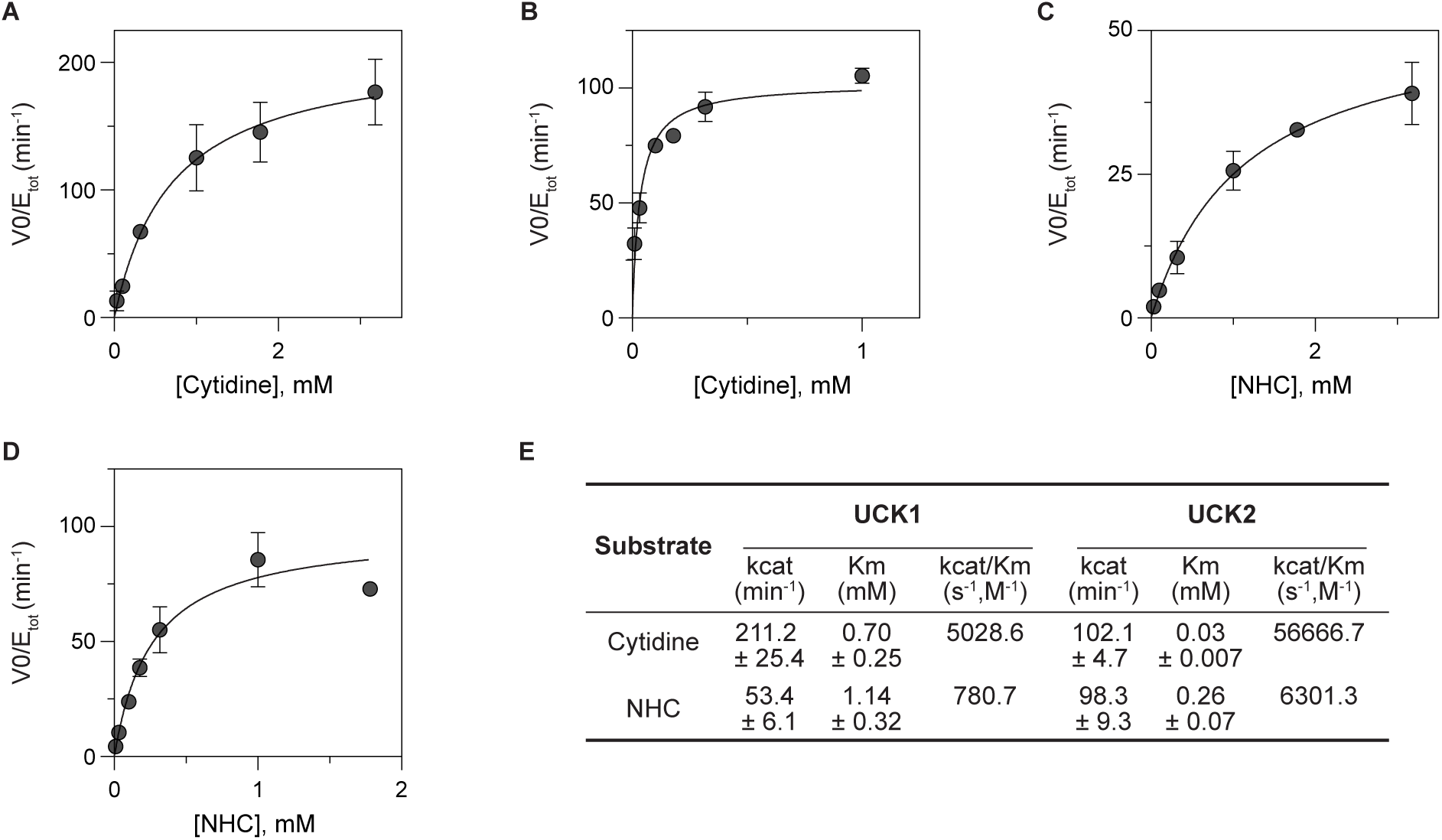
UCK2 is the preferred kinase in phosphorylating molnupiravir metabolite NHC. **A-B.** Saturation curves of UCK1 (A) or UCK2 (B) -mediated phosphorylation of cytidine, using ADP-coupled activity assay. **C-D.** Saturation curves of UCK1 (C) or UCK2 (D) -mediated phosphorylation of NHC, using ADP-coupled activity assay. In A-D, mean initial rates ± SEM of n=2 independent experiments performed in triplicates are shown. **E.** Kinetic parameters of UCK1 and UCK2-mediated phosphorylation of NHC, in comparison to cytidine, a canonical substrate of UCK1 and UCK2. Kinetic parameters ± SEM were determined via Michaelis-Menten equation (GraphPad Prism) based on the saturation curve shown in A-D.

### UCK2 downregulation hampered the anti-SARS-CoV-2 efficacy of NHC by limiting its bioactivation into NHC-TP

As the more efficient kinase UCK2 demonstrated moderate activity against NHC compared to its physiological substrate cytidine, we next examined if the activity of UCK2 could be pharmacologically relevant for NHC efficacy in cells. We then utilized a previously established high-throughput-compatible infectivity assay setup [26] to interrogate the the antiviral activity of NHC against SARS-CoV-2, its clinical indication. Specifically, A549/ACE2 cells, a coronavirus susceptible lung basal epithelial cell line overexpressing coronavirus receptor ACE2, were infected with a patient-derived SARS-CoV-2 Wuhan-Hu-1 strain (Pango lineage B; GenBank: MT093571). Viral infectivity was subsequently determined based on the percentage of infected cells as indicated by positive immunofluorescent staining of SARS-Co-V-2 nucleocapsid protein (**Fig. 4a**, **Fig. S4a-b**). The platform utilized high-content imaging analysis and automated CellProfiler pipeline for speedy image acquisition and analysis, respectively, thereby accommodating over 3,000 cells per condition and drug dose-response matrix setup. Overall, the assay setup greatly improved experiment throughput and data reproducibility compared to traditional immunofluorescence-based viral quantification methods, e.g. focus-forming assay [27].

**Figure 4.**
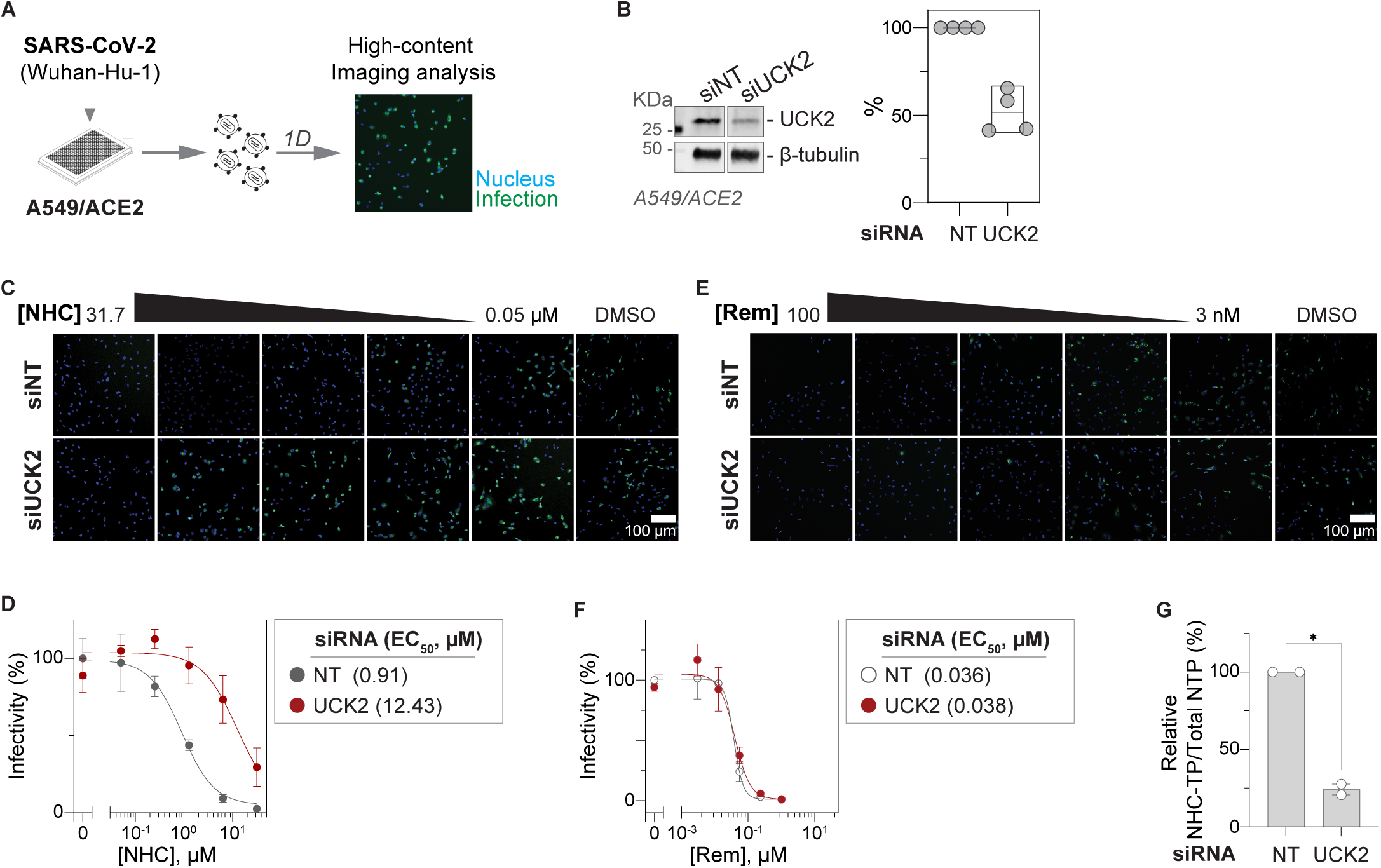
UCK2 expression dictates the anti-SARS-CoV-2 efficacy of NHC. **A.** Schematic representation of the high content imaging-based infectivity assay setup. Lung alveolar basal epithelial A549/ACE2 cells, engineered to overexpress SARS-CoV receptor ACE2, were infected with SARS-CoV-2 at a MOI of 0.07 overnight, before viral infection was assayed via immunofluorescence staining against the viral nucleocapsid protein (NC) and cell nucleus (DAPI). Infectivity was determined as percentage of DAPI/NC double-positive cells out of DAPI single-positive cells, which was further normalized to infectivity level in virus-infected wildtype group and presented as relative infectivity %. **B**. Western blot analysis of A549/ACE2 cell lysates 72 hours post-transfection with UCK2-targeting (siUCK2) or control (siNT) siRNAs. *Left panel*: Representative Western blot image. *Right panel*: densitometry analysis of n = 4 independent experiments, where protein signals were normalized to β-tubulin and relative to that of siNT-treated cells. **C-F.** siRNA-mediated downregulation of UCK2 in A549/ACE2 cells antagonized the anti-SARS-CoV-2 efficacy of NHC (C-D), but not another antiviral nucleoside analogue remdesivir (Rem, E-F). At 72 hours post-transfection with UCK2-targeting (siUCK2) or control (siNT) siRNAs, A549/ACE2 cells were re-seeded and infected with SARS-CoV-2 overnight in the presence of antivirals or the drug diluent control DMSO, before viral infectivity was determined using the high content imaging-based infectivity assay as depicted in A. In C and E, representative immunofluorescence staining images of infected A549/ACE2 cells, scale bars represent 100 μm; in D and F, mean relative infectivity % ± SEM of n=3 independent experiments performed in duplicate are shown. **G.** UCK2 knockdown antagonized NHC through limiting the intracellular levels of its tri-phosphorylated active metabolite NHC-TP. A549/ACE2 cells transfected with UCK2-targeting (siUCK2) or control (siNT) siRNAs were treated with 100 µM NHC for 6 hours before cell lysates were harvested for the measurement of intracellular NHC-TP levels. The NHC-TP levels were normalized to total NTP levels and then to siNT samples. Mean of the resulting relative NHC-TP levels ± SEM of n=2 independent experiments are shown. Paired t-test was performed across treatment groups [Relative NHC-TP (siNT) vs. Relative NHC-TP (siUCK2), p = 0.03, t = 21.6, df = 1], where asterisks signify statistical significance (∗p ≤ 0.05).

To delineate the role of UCK2 in the antiviral efficacy of NHC, we next conducted the SARS-CoV-2 infectivity assay using A549/ACE2 cells transfected with a pool of four UCK2-specific siRNAs (siUCK2) or non-targeting control siRNA (siNT), done in the presence or absence of NHC or another SARS-CoV-2 drug remdesivir. Both being NA drugs, NHC and remdesivir share common antiviral mechanism, i.e. target viral RNA polymerase for misincorporation into viral genome upon tri-phosphorylation. Nevertheless, remdesivir is a purine analogue and requires adenylate kinase AK2 for bioactivation [28], thereby serving as the control antiviral drug to decipher potential roles of UCK on NHC efficacy - via the bioactivation versus downstream pathways/targets of NHC. In A549/ACE2 cells with endogenous UCK2 levels, NHC or remdesivir demonstrated micromolar and high nanomolar antiviral EC50 values, respectively, agreeing with previous studies [29–31] (**Fig. 4b-d, Fig. S4a-b**). Upon UCK2 knockdown via siUCK2 transfection, antiviral efficacy of NHC was significantly hampered as exemplified by a drastic increase of the antiviral EC50 value by over 10X, which was not observed for the control drug remdesivir (**Fig. 4c-d**). Of note, during the course of the infectivity assay, UCK2 KD alone or in combination with either antiviral, had no adverse effect on the host cell proliferation/viability (**Fig. S4c-d**).

These data strongly suggest that UCK2 KD antagonized NHC by hampering its bioactivation, which was further supported by determining the intracellular pool of NHC-TP, the antiviral metabolite of NHC. A549/ACE2 cells were transfected with siNT/UCK2 siRNA, and one day post-transfection, cells were treated with NHC followed by intracellular nucleotide extraction and measurement via strong anion-exchange HPLC. Whilst endogenous NTP levels were minimally affected, UCK2 KD significantly reduced the intracellular pool of NHC-TP by 75% (**Fig. 4g, Fig. S4f-g**), directly highlighting the pivotal role of UCK2 in the bioactivation and thereafter antiviral efficacy of NHC.

### UCK is a viable target for tailoring antiviral efficacy of NHC

Given the critical role of UCK in dictating the therapeutic efficacy of NHC/molnupiravir, we next sought to investigate its potential as a pharmacological target for tailoring the drug response. We first investigated if the antiviral efficacy of NHC could be modulated via regulating the catalytic activity of UCK, by employing the cell-active UCK inhibitor CPU. Specifically, A549/ACE2 cells of wildtype UCK expression were infected with SARS-CoV- 2 in the presence of increasing concentrations of CPU in combination with NHC or the control antiviral remdesivir, before viral infectivity was determined using the immunofluorescence staining-based infectivity assay. Without inhibiting host cell proliferation/viability, the addition of the UCK inhibitor CPU significantly antagonized the anti-SARS-CoV2 efficacy of NHC in a dose-dependent manner, elevating the antiviral EC50 value by up to 7X fold compared to the DMSO control group (**Fig. 5a-b, Fig. S5a-c**). Concurrently, direct measurement of the intracellular NTP species revealed an 80% reduction of NHC-TP pool, upon the addition of CPU (**Fig. 5c, Fig. S5d-e**). Meanwhile, no antagonism was observed for the control antiviral remdesivir (**Fig. 5d-e, Fig. S5f**).

**Figure 5.**
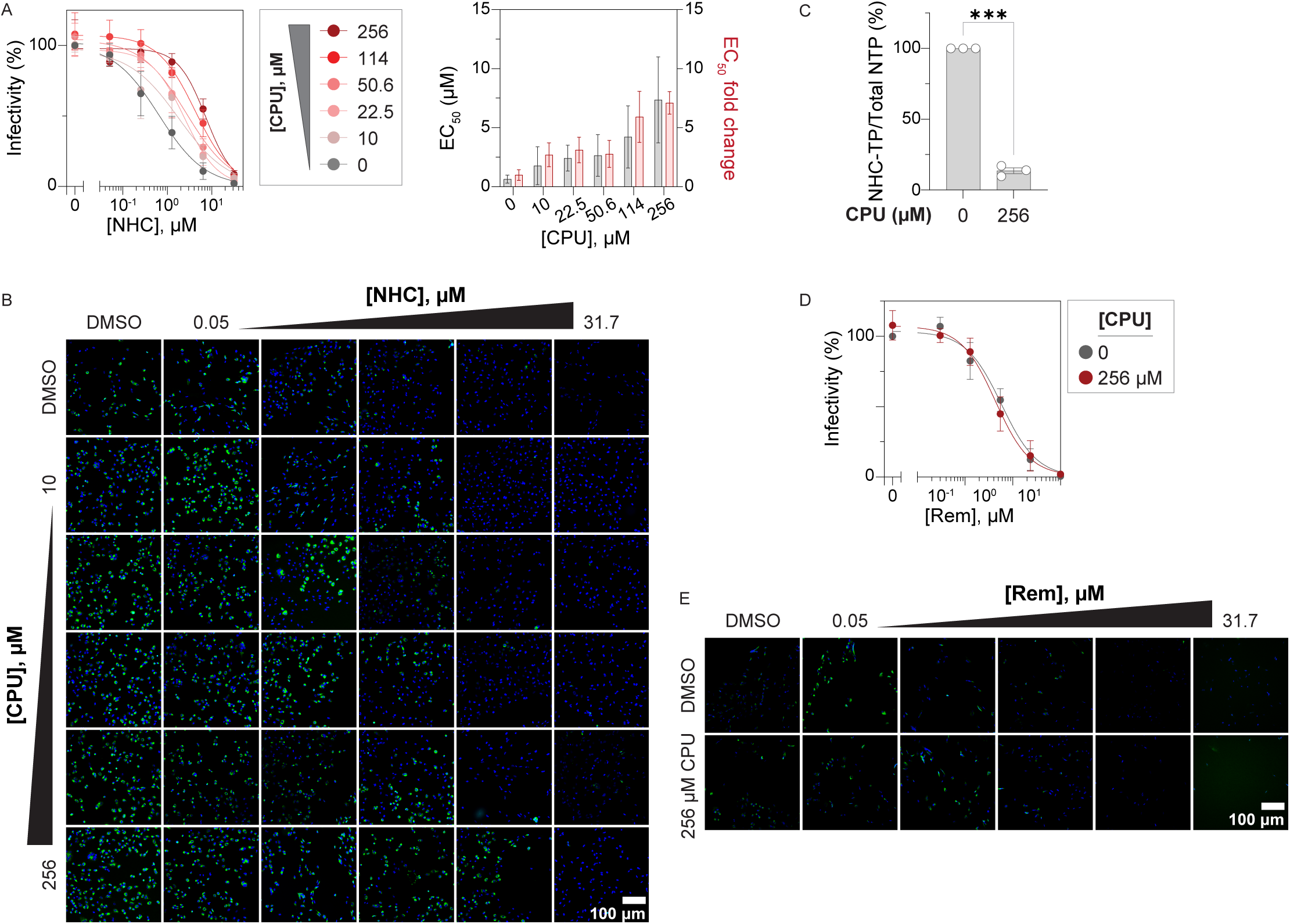
UCK inhibition antagonizes the anti-SARS-CoV-2 efficacy of NHC. **A-D.** CPU-mediated UCK inhibition in A549/ACE2 cells dose-dependently antagonized the anti-SARS-CoV-2 efficacy of NHC, but not another antiviral nucleoside analogue remdesivir. A549/ACE2 cells were infected with SARS-CoV-2 overnight in the presence of dose-response matrix composed of the antivirals/DMSO and CPU/DMSO, before viral infectivity was determined using the high content imaging-based infectivity assay as depicted in Fig. 4 A. In B (NHC) and D (remdesivir, Rem), representative immunofluorescence staining images of infected A549/ACE2 cells, scale bars represent 100 μm. In A (left panel, NHC) and C (remdesivir, Rem), mean relative infectivity % ± SEM of n=4 independent experiments performed in duplicate are shown. The antiviral EC_50_ (half maximal effective concentration) of NHC with increasing concentration of CPU and the EC_50_ fold changes relative to DMSO control group are further shown in A (right panel). **E.** UCK2 inhibition by CPU significantly hampered the accumulation of intracellular NHC-TP. A549/ACE2 cells were treated with DMSO or 256 µM CPU for 1 hour before cells were further treated with 100 µM NHC for another 5 hours. Cell lysates were subsequently harvested, and the intracellular nucleotide levels were determined. The NHC-TP levels were normalized to total NTP levels and then to DMSO control samples. Mean of the resulting relative NHC-TP levels ± SEM of n=3 independent experiments are shown. Paired t-test was performed across treatment groups [Relative NHC-TP (0 µM CPU) vs. Relative NHC-TP (256 µM CPU), p = 0.0006, t = 40.51, df = 2], where asterisks signify statistical significance (∗∗∗p ≤ 0.001).

A high selectivity index (SI) – defined as the ratio of cell growth inhibitory EC50 to the antiviral EC50, entails potent antiviral activity with minimal cellular toxicity, and potentially successful clinical responses [19]. In the prolonged 4-day cell proliferation assay, NHC dose-dependently inhibited proliferation of multiple cell lines (**Fig. 1**), accompanied by aberrant cell cycle profile, i.e. collapse of replicating S-phase concurrent to G1 arrest (**Fig. S6a-b**), indicating that NHC has off-target cytotoxicity potentially via interfering with genome replication. As UCK inhibition via CPU could antagonize both the cytotoxicity as well as the antiviral efficacy of NHC (**Fig. 5a-b**), we then determined if UCK could govern the selectivity index of NHC/molnupiravir, specifically using growth inhibitory EC50 values from 4-day resazurin assay and antiviral EC50 values. Critically, in A549/ACE2 cells, both UCK expression downregulation via siUCK2, as well as UCK activity inhibition via CPU, significantly reduced selectivity index of NHC by up to 2X fold (**Fig. 6**, **Fig. S6c**), overall highlighting that UCK is a viable and critical target for achieving optimal antiviral efficacy, and furthermore therapeutic window of the anti-SARS-CoV-2 drug molnupiravir.

**Figure 6.**
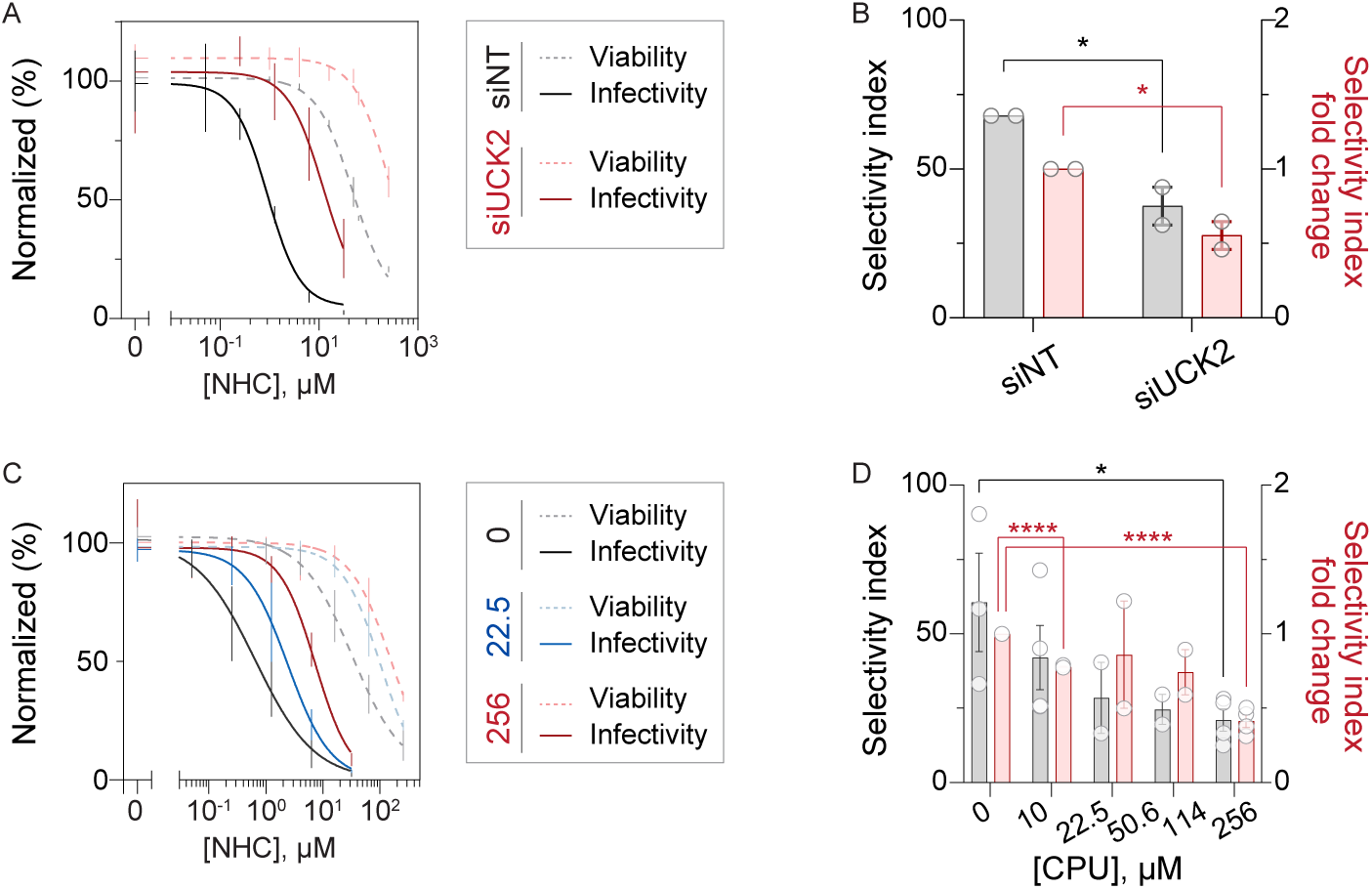
UCK is critical for an optimal selectivity index of NHC. **A-B.** Depletion of UCK2 significantly reduced the selectivity index of NHC by 50%. A549/ACE2 cells transfected with UCK2-targeting (siUCK2) or control (siNT) siRNAs were treated with NHC for 4 days before cell viability was determined using resazurin reduction assay. Alternatively, cells were infected with SARS-CoV-2 overnight in the presence of NHC before viral infectivity was determined using high content imaging-based infectivity assay. Mean viability/infectivity relative to DMSO group ± SEM of n=2 independent experiments are shown in A, and were further curve fitted using a nonlinear curve fitting model with variable slope (GraphPad Prism) to generate the antiviral and growth inhibitory EC_50_ values of NHC. Selectivity index was then calculated as the ratio of growth inhibitory EC_50_ and antiviral EC_50_. In B, mean selectivity index ± SEM, and mean fold change of selectivity index ± SEM are shown. Student’s t-tests were performed across treatment groups [Selectivity index (siNT) vs. selectivity index (siUCK2), p = 0.041; selectivity index fold change (siNT) vs. selectivity index fold change (siUCK2), p = 0.041)], where asterisks signify statistical significance (∗p ≤ 0.05). **C-D.** Inhibition of UCK activity by CPU dose-dependently reduced the selectivity index of NHC, by up to over 50%. A549/ACE2 cells were treated with a dose-response matrix of CPU and NHC for 4 days before cell viability was determined using resazurin reduction assay, or alternatively, cells were infected with SARS-CoV-2 overnight before viral infectivity was determined. Mean viability/infectivity relative to DMSO group ± SEM of n=2-4 independent experiments are shown in C, and were further curve fitted using a nonlinear curve fitting model with variable slope (GraphPad Prism) to generate the antiviral and growth inhibitory EC_50_ values of NHC. Selectivity index was then calculated as the ratio of growth inhibitory EC_50_ and antiviral EC_50_. In D, mean selectivity index ± SEM, and mean fold change of selectivity index ± SEM are shown. Student’s t-tests were performed across treatment groups [Selectivity index (0 µM CPU) vs. selectivity index (10 µM CPU), p = 0.368; selectivity index (0 µM CPU) vs. selectivity index (22.5 µM CPU), p = 0.259; selectivity index (0 µM CPU) vs. selectivity index (50.6 µM CPU), p = 0.195; selectivity index (0 µM CPU) vs. selectivity index (256 µM CPU), p = 0.042; selectivity index fold change (0 µM CPU) vs. selectivity index fold change (10 µM CPU), p = 1.04E-7; selectivity index fold change (0 µM CPU) vs. selectivity index fold change (22.5 µM CPU), p = 0.636; selectivity index fold change (0 µM CPU) vs. selectivity index fold change (50.6 µM CPU), p = 0.107; selectivity index fold change (0 µM CPU) vs. selectivity index fold change (256 µM CPU), p = 8.14E-5], where asterisks signify statistical significance (∗p ≤ 0.05, ∗∗∗∗p ≤ 0.0001).

## DISCUSSION

Molnupiravir is a broad-spectrum NA antiviral drug against several pathologically important RNA viruses and has been used clinically to treat SARS-CoV-2 infection. Whilst its antiviral mechanism-of-action has been thoroughly studied and deciphered, pharmacogenetic host factors that regulate its therapeutic responses – efficacy as well as selectivity – have not been mechanistically defined and characterized. Here in this study, we identified that host pyrimidine salvage enzymes UCK1 and UCK2 catalyze the first phosphorylation and bioactivation step for NHC – the active compound of molnupiravir – and thereby dictate its antiviral efficacy and more critically, therapeutic window.

In target engagement and enzyme kinetic studies using recombinant enzymes, UCK1 and UCK2 were able to directly bind to NHC and catalyze its phosphorylation (**Fig. 2-3**). Subsequently, using a SARS-CoV-2 infectivity model, downregulation of host cell UCK2 expression via siRNA transfection led to NHC resistance (**Fig. 4**) exemplified by a drastic reduction of intracellular NHC-TP – the active metabolite of molnupiravir – and consequently a 10X increase of the drug antiviral EC50. Similar to our data with NHC, UCKs have been demonstrated to phosphorylate and thereby activate other NA drugs, such as azacitidine [1]. The principal mechanism of azacitidine resistance has been attributed to loss-of-activity mutations of UCK2 and loss-of-expression of UCK1, in leukemic cell line models [7, 9, 32] and patients [6], respectively. This underscores that UCKs could be biomarkers informing treatment outcome for NA drugs, and our data extends this to molnupiravir/NHC. Whilst few catalytically dead UCK2 variants have been identified in leukemic cell lines [9], no studies have been conducted to recapitulate these findings in patient material, or to identify such variants for UCK1 [6]. To fully decipher the potential of UCK1 and UCK2 as clinically applicable biomarkers for NA therapy including molnupiravir/NHC, we therefore envision that patient variant profiling of UCKs and subsequent correlation to drug response should be the focus of future studies.

As UCKs phosphorylated NHC and further controlled its antiviral efficacy, we next investigated if they are pharmacologically tractable targets for tailoring NHC response. To that end, we employed a pan-UCK inhibitor CPU in the SARS-CoV-2 infectivity assay, and observed that CPU dose-dependently antagonized the antiviral efficacy of NHC (**Fig. 5**). More interestingly, with both CPU treatment and UCK2 knockdown, the hampered efficacy is accompanied by a narrowed selectivity index of NHC, collectively suggesting that elevated UCK expression/activity promotes optimal drug efficacy and therapeutic window (**Fig. 6**). NHC, upon being converted to NHC-TP, is used by the viral polymerase in place of CTP and is then ambiguously matched with GTP or ATP in the subsequent replication cycles, leading to a lethal mutation burden in the viral genome and generation of infection-defective progeny virions [16]. Similarly, a recent study showed that NHC could also induce these mutations in cellular though non-human genome, as illustrated using mouse lymphoma and myeloid cell lines [18]. The study further implicated UCK2 as the principal driver for such mutations [18]. Nevertheless, the study did not mechanically characterize the phenotypic consequences of these mutations, in cell cycle progression, cell viability, and/or cell proliferation, making the role of UCK2 in NHC-induced cytotoxicity inconclusive. Here in our study, when using CPU at concentrations where antagonism of antiviral efficacy is consistently observed (e.g. 50-256 µM), minimum rescue of the NHC-induced cytotoxicity was achieved, suggesting that NHC can induce cytotoxicity through additional pathway(s)/drug metabolites aside from those activated via UCKs (**Fig. S6C**). Systematic profiling of host metabolic factors governing NHC/monulpiravir efficacy and selectivity could potentially provide additional targets, alone or in combination with UCKs, for achieving optimal drug responses against its clinical indication SARS-CoV-2, as well as other pathologically important RNA viruses.

UCKs have two isoforms of different catalytic efficiencies, with UCK2 often displaying superior activities compared to UCK1, for the canonical substrates cytidine and uridine as well as NA drugs [1]. Here, in kinetic studies using recombinant UCKs, UCK2 showed markedly higher activities towards NHC as compared to UCK1, displaying a 9-fold higher catalytic efficiency (**Fig. 3**). Nevertheless, UCK2 expression is often limited to embryo and placenta and only upregulated in tissues upon malignant transformation, whilst UCK1 is more ubiquitously expressed, relevant to the multiorgan tropism of coronavirus infection [33]. Our kinetic studies revealed that, compared to UCK1, UCK2 had 5 times lower Km –0.26 ± 0.07 mM (UCK2) Vs. 1.14 ± 0.32 mM (UCK1), and only twice higher kcat – 98.3 ± 9.3 min^-1^ (UCK2) Vs. 53.4 ± 6.1 min^-1^ (UCK1), suggesting its higher catalytic efficiency towards NHC was mainly attributed to the higher substrate binding affinity. With sustained and higher intracellular accumulation of NHC and metabolites compared to the drug level in plasma [34, 35], the inferior affinity of UCK1 towards NHC could potentially be overcome *in vivo*, making UCK1 equally relevant to NHC bioactivation. Established cell lines however are often of cancerous/transformed origins, and thus express high levels of UCK2 [36] and potentially have metabolic re-wiring [2], collectively confounding the role(s) of UCK1. Hence, to fully evaluate and decipher the involvement of UCK1 in controlling NHC/molnupiravir efficacy, we envision future studies employing patient materials/data are warranted.

In summary, our study establishes the pivotal role of UCKs in dictating molnupiravir therapeutic efficacy and highlights these enzymes as viable targets for tailoring its response, providing further incentive to develop UCK probes – inhibitory and activating – which is currently an understudies area of research.

## ACKNOWLEDGEMENTS

We thank the Protein Science Facility (http://ki.se/psf) and BSL3 Biomedicum Core Facility at Karolinska Institutet for recombinant protein production and infectivity assay setup, respectively. We thank Professor Anna Karlsson (Karolinska Institutet) for kindly providing the pET-15b-UCK1 expression plasmid.

The authors acknowledge funding from the Swedish Research Council (2018-02114 to S.G.R.), Swedish Cancer Society (19-0056-JIA, 20-0879-Pj, and 23-2782-Pj to S.G.R.), the Swedish Children’s Cancer Fund (PR2019-0014 and PR2022-0003 to S.G.R.), Karolinska Institutet in the form of a Board of Research Faculty Funded Career Position (to S.G.R.), the Felix Mindus contribution to Leukemia Research (2020-02573 and 2022-02835 to S.M.Z.), the Karolinska Institute foundation for virus research (2020-00249 and 2022-00247 to S.M.Z.), Loo and Hans Osterman Foundation for Medical Research (2022-01262 to S.M.Z.), Åke Wibergs Foundation (M22-0011 and M23-0088 to S.M.Z.), Stiftelsen Lars Hiertas Minne (F0222-0147 to S.M.Z.), Stiftelsen Clas Groschinskys Minnesfond (M2353 and M2449 to S.M.Z.), and Karolinska Institutet Research Grants (2024-02905 to S.M.Z.)

## AUTHOR CONTRIBUTIONS

H.S., S.A., M.T., N.C.K.V., M.A. and S.M.Z. designed, performed, and/or analyzed biological experiments; H.S., S.S., L.F., A.B.P.K., A.C. and S.M.Z. designed, performed, and/or analyzed biochemical experiments; S.M.Z. conceptualized the study, and further compiled data and prepared the manuscript with H.S. and S.G.R.; S.G.R and S.M.Z. supervised the project. All authors discussed the results and approved the manuscript.

## METHODS AND MATERIALS

### Resource availability

#### Lead contact

Further information and requests for resources and reagents should be directed to and will be fulfilled by the lead contact, Si Min Zhang.

#### Materials availability

Materials and compounds generated in this study are available from the corresponding authors on reasonable request.

#### Data and code availability

Original western blot images have been deposited at Mendeley and are publicly available as of the date of publication.

### Experimental model and study participant details

#### Reagents

*Antibodies* – The antibody against SARS-CoV-2 nucleocapsid protein (mouse, cat. No. MA1-7404) was purchased from Invitrogen. Antibodies against UCK1 (rabbit, cat. No. 12271-1-AP) and UCK2 (rabbit, cat. No. 10511-1-AP) were purchased from Proteintech. Antibody against β-actin (rabbit, cat. no. ab6046) was purchased from Abcam. Donkey anti-rabbit IRDye® 800CW (cat. No. 926-32213) was purchased from Li-Cor. Alexa Fluor™ 488-conjugated goat anti-mouse IgG (H+L) cross-adsorbed secondary antibody (cat. No. A21202) was purchased from Invitrogen.

*Chemicals* – Cyclopentenyl uracil (CPU, cat. No. SML2972-5MG), cytarabine (cat. No. C1768), gemcitabine (cat no. G6423), 5’-azacitidine (cat. No. A2385), cytidine (cat. No. C4654-5G), thymidine (cat. No. T9250), and ATP (cat. No. A2383-1G) were purchased from Sigma-Aldrich; cyclopentenyl cytosine (CPC, cat. No. NSC 375575) was acquired from NCI; 3’-deazauridine (DAU, cat. No. ND16524) was purchased from Carbosynth; molnupiravir (cat. No. HY-135853) and NHC (cat. No. HY-125033) were purchased from MedChemExpress; idoxuridine was purchased from Fluka Biochemika; decitabine (cat. No. QB-5169) was purchased from Combi-Blocks; and floxuridine was purchased from Ega-Chemie. ATP was dissolved in dH2O at 50 mM and stored at -20 °C, and other chemicals were dissolved in DMSO and stored at 4 °C.

*Recombinant protein* – Recombinant UCK1 with a N-terminal His tag was expressed and purified by Protein Science Core Facility (PSF), Karolinska Institutet, Sweden; and recombinant UCK2 with a C-terminal His tag (cat. No. NBP1-50888-0.1mg) was obtained from Novus Biologicals. The expression plasmid pET-15b-UCK1 encoding a human UCK1 construct with N-terminal His-tag was kindly provided by Professor Anna Karlsson (Karolinska Institutet, Sweden)[1]. Briefly, the expression plasmid pET-15b-UCK1 was transformed into *E.coli* BL21 (DE3) T1R pRARE2 cells. The cells were cultivated in Terrific Broth (TB) medium and protein expression was induced by 0.5mM isopropyl-β-D-1-thiogalactopyranoside (IPTG). Cells were then harvested by centrifugation and resuspended in IMAC lysis buffer supplemented with cOmplete Mini, EDTA-free protease inhibitor (Roche), and then stored at -80°C. During protein purification, cell pellets were thawed with benzonase (2.5 U/mL, Merck-Millipore) and then disrupted by pulsed sonication (4s/4s 4 min, 80% amplitude), before lysates were subject to centrifugation (20 min at 49000 × g) followed by filtering through 0.45 μm filters. Protein was then purified from the clarified lysates firstly on a HisTrap column (GE Healthcare), using buffer A (20 mM HEPES, 500 mM NaCl, 10% glycerol, 10 mM imidazole, 0.5 mM TCEP, pH 7.5) as starting buffer and an imidazole gradient (10–500 mM) in buffer A as the elution buffer. Protein was then further purified using a Gel Filtration Column, HiLoad 16/60 Superdex 200 (GE Healthcare) in 20 mM HEPES, 300 mM NaCl, 10% glycerol, 0.5 mM TCEP, pH 7.5. Protein-containing fractions were confirmed by SDS-PAGE, pooled in storage buffer (20 mM HEPES, 300 mM NaCl, 10% glycerol, 2 mM TCEP, pH 7.5), aliquoted and stored at 80°C. Protein purities were confirmed using SDS-PAGE and Coomassie staining (see **Fig. S7**), and concentrations were determined by NanoDrop (Thermo Fisher Scientific) A280 measurement.

#### Biological Resources

*Cell lines –* LCL-889 cells were obtained by immortalizing healthy donor B cells, as described previously [37]. HL-60 (cat. No. CCL-240), THP-1 (cat. No. TIB-202), MV4-11 (cat. No. CRL-9591), KG1a (cat. No. CCL-246.1), MOLT-4 (cat. No. CRL-1582), CCRF-CEM (cat. No. CCL-119), and DAUDI (cat. No. CCL-213) were acquired from ATCC. Ramos, BL-41, Raji, MOLT-16, and Wil2-NS cells were acquired from DSMZ. PL-21 was kindly gifted by Dr. Sören Lehmann (Karolinska Institutet, Sweden); DAOY was kindly gifted by Dr. Fredrik Johansson Swartling (Uppsala Universitet, Sweden); UW228-3 was kindly gifted by Dr. John Inge Johnsen (Karolinska Institutet, Sweden); U-343 MG and U-251 MG were kindly gifted by Dr. Lars Braeutigam (Karolinska Institutet, Sweden); and LN-229 was kindly gifted by Dr. Francoise Dantzer (University of Strasbourg, France); A549/ACE2 and Vero E6 were kindly gifted by BSL3 Biomedicum Core Facility (Karolinska Institutet, Sweden).

THP-1, HL-60, and KG1a were cultured in IMDM medium (cat. No. 12440061); U-251 MG, LN-229, and Vero E6 were cultured in DMEM medium with high glucose and GlutaMAX ^TM^ (cat. No. 31966047); DAOY and U343 were cultured in MEM medium with GlutaMAX^TM^ (cat. No. 41090036), supplemented with 1mM pyruvate and 1X NEAA (cat. No. 11350912); UW228-3 was cultured in DMEM/F12 medium with Glutamax^TM^ (cat. No. 10565018); A549/ACE2 was cultured in DMEM/F12 medium with Glutamax^TM^ supplemented with 1X NEAA; and all the other cell lines were cultured in RPMI 1640 with GlutaMAX™ (cat. No. 61870044), at 37°C with 5% CO2 in a humidified incubator. All the cell lines were cultured in medium supplemented with 10% heat-inactivated fetal bovine serum (cat. No. 10500064) and penicillin/streptomycin (100 U/mL and 100 µg/mL, respectively, cat. No. 15070063), except for KG1a and PL-21, which were cultured in 20% fetal bovine serum and penicillin/streptomycin (100 U/mL and 100 µg/mL, respectively). All cell culture reagents were purchased from ThermoFisher Scientific. The cell lines were regularly monitored and tested negative for the presence of mycoplasma using a commercial biochemical test (MycoAlert, Lonza).

*Virus –* The clinical isolate SARS-CoV-2 (Pango lineage B; GenBank: MT093571) was propagated in African green monkey kidney epithelial cells Vero E6 cells. Briefly, viral inoculated cells were cultured in DMEM medium supplemented with 5% fetal bovine serum and penicillin/streptomycin (100 U/mL and 100 µg/mL, respectively) for 72 hours at 37°C with 5% CO2 in a humidified incubator, till cytopathic effect was apparent. Cells were subsequently harvested, and supernatant was recovered via centrifugation at 2000 rpm for 5 minutes. Clarified supernatant was then aliquoted and stored at -80 °C. Viral titre was determined using plaque assay. Vero E6 cells at 80% confluency in a 6-well plate were inoculated with serial diluted virus aliquots in DMEM medium at 37 °C for 1 hour. Subsequently, cells were washed by PBS twice and overlaid with 0.4% carboxymethylcellulose in MEM supplemented with 5% fetal bovine serum and penicillin/streptomycin (100 U/mL and 100 µg/mL, respectively) at 1mL/well, followed by incubation for 48 hours at 37°C with 5% CO2 in a humidified incubator, till plaques were visible. Cells were then fixed with 10% formaldehyde for 1 hour at room temperature, followed by washing with ddH_2_O thrice and incubation with crystal violet for 30 minutes at room temperature. Following washing in ddH_2_O and air-drying, plaques were then counted and used to estimate the viral titre following the formula 𝑉𝑖𝑟𝑎𝑙 𝑡𝑖𝑡𝑒𝑟 = 𝐴𝑣𝑔_# 𝑝𝑙𝑎𝑞𝑢𝑒𝑠_/(𝐷𝑖𝑙𝑢𝑡𝑖𝑜𝑛 𝑓𝑎𝑐𝑡𝑜𝑟 × 𝐼𝑛𝑜𝑐𝑢𝑙𝑢𝑚 𝑣𝑜𝑙𝑢𝑚𝑒).

*Bacteria -* E. coli BL21(DE3) was obtained from Invitrogen.

### Method details

#### In silico UCK substrate screening library composition

In silico substrate screening was performed on a curated library composed of 3323 therapeutics, sourced from SelleckChem (Catalog No. L7200) and pharmacopeia (Catalog No. L1300). Compound information including (1) name, (2) formula, (3) CAS, and (4) coordinates, was extracted and was subsequently used to identify potential substrates for UCKs, based on structural similarity to the canonical substrates of UCKs, i.e. cytidine and uridine.

### Cell viability assays

#### Resazurin reduction viability assay

The resazurin reduction viability assay was conducted as described previously [38, 39]. Briefly, suspension and adherent cells were seeded into 384-well or 96-well assay plates at 50,000 /mL or 25,000/cm^2^, respectively; and compounds of indicated concentrations were added using a D300e Digital Dispenser (Tecan). When applicable, DMSO levels were normalised across the plate at the maximum level of 1%. Suspension cells were treated on the same day of cell seeding and adherent cells were treated one day post-seeding. Cells were then incubated at 37°C with 5% CO2 in a humidified incubator for 1 or 4 days. On the day of assaying, cells were incubated with resazurin sodium salt (10 µg/mL) for 4-6 hours and the reduction of resazurin by viable cells were subsequently assessed by measuring fluorescence intensity at 544/590 nm (Ex/Em) using a HidexSense plate reader (Hidex). Relative cell viabilities were calculated by subtracting signals of medium-only negative control wells, and then normalized to signals of cell-only positive control wells. When applicable, relative cell viabilities were used to generate compound EC_50_ values *via* a nonlinear curve fitting model with variable slope (GraphPad Prism), as well as to calculate drug combination synergy scores using the Bliss model in SynergyFinder (https://synergyfinder.fimm.fi/) [40].

#### Cell growth assessment via cell confluency

Growth of A549/ACE2 cells of indicated treatment was assessed via cell confluency readout module of Tecan Spark Cyto, and cell confluency % was determined by the machine inbuilt analysis module.

#### Western blot

Cells were harvested by centrifugation at 400×g for 4 minutes or trypsinization followed by centrifugation, and were subsequently lysed on ice for 30 minutes using RIPA buffer supplemented with cOmplete Mini, EDTA-free protease inhibitor (Roche) and Halt^TM^ phosphatase inhibitor (Thermo Fisher Scientific). Lysates were then clarified via centrifugation at 12000×g for 20 minutes, followed by protein concentration determination using Pierce ^TM^ BCA protein assay kit (Thermo Scientific), per manufacturer’s instruction. For Western blot, clarified cell lysates were mixed in β-mercaptoethanol-supplemented Laemmli buffer (Bio-Rad) and then heated at 99°C for 5-10 min. Lysates containing 15-20 µg protein were subject to sodium dodecyl sulfate-polyacrylamide gel electrophoresis (SDS-PAGE) using 4–15% Mini-PROTEAN TGX gels, and proteins were subsequently transferred to nitrocellulose membranes using a Trans-Blot Turbo machine (Bio-Rad). Following blocking with Odyssey Blocking Buffer (LI-COR), membranes were first probed with primary antibodies at RT for 1 h or 4°C overnight, and then probed with species-appropriate secondary antibodies at RT for 30 min. Between incubations, membranes were washed trice using TBST (Tris-buffered saline, 0.1% Tween 20). Protein bands were visualised using an Odyssey Fc Imaging System (Li-Cor Biosciences), and subsequently analysed using Image Studio Lite (Ver. 5.2, Li-Cor Biosciences). All uncropped western blot images are provided in the source data.

#### Differential scanning fluorometry (DSF)

DSF assay on recombinant UCK was conducted as described previously, with minor modifications [38]. Briefly, in Axygen 384-well PCR microplate, 5 µM recombinant UCK proteins in 20 µL assay buffer (50 mM Tris-HCl (pH = 7.5), 5 mM MgCl_2_, 100 mM KCl, 2 mM DTT) fortified with Sypro Orange (10X, Invitrogen), were mixed with 5mM ATP, alone or in the presence of compounds of indicated concentrations. The assay mixture was then subject to a 20-85/90°C temperature gradient for 20 min, with the fluorescence intensities (RFU) measured every second using a LightCycler 480 Instrument II (Roche Life Science). Melting curves and negative derivative (-*d*RFU/*d*T) transformation were subsequently generated, and melting temperatures (Tm) were identified as the local minima of the negative derivative curves, done using the machine inbuilt software LightCycler 480 SW 1.5.1.

#### ADP-Glo coupled UCK kinase activity assay

UCK kinase activity was determined by coupling the kinase reaction to ADP-Glo™ kinase assay (Promega). In 384-well plates, recombinant UCK protein in 5µL reaction buffer (50 mM Tris-HCl, pH = 7.5, 5 mM MgCl2, 100 mM KCl, 2 mM DTT, 0.5 mg/mL BSA) was mixed with ATP at a final concentration of 50 µM, followed by the addition of substrates at indicated concentrations to initiate the reaction. For enzyme titration, substrate final concentration was 100µM for both cytidine and NHC; and the final concentration of UCK1 ranged between 0.00625 to 1.6 µM for cytidine and 0.1 to 1.6 µM for NHC, and UCK2 ranged between 0.938 to 15 nM for cytidine and 3.75 to 60 nM for NHC. For kinetic studies, the final concentration of UCK1 was 6.25 nM for cytidine and 41.6 nM for NHC, and UCK2 of 0.9375 nM for cytidine and 3.75 nM for NHC.

Reactions then proceeded up to 100 min, which were terminated with 8-20 min intervals with the addition of 5 μL ADP-Glo reagent (ATP depletion reagent) followed by incubation at room temperature for 1 hour. Subsequently, 10 μL kinase detection reagent was added followed by 1 hour incubation at room temperature in the dark. Subsequently, luminescence was detected on a HidexSense plate reader (Hidex). ADP formation was calculated using an ADP/ATP conversion standard curve, and was further used to estimate the initial reaction rates and subsequently, kinetic parameters, via Michaelis-Menten equation (GraphPad Prism).

#### RNA interference transfection

Transfections were performed using INTERFERin (Polyplus Transfection) following manufacturer’s instructions. UCK2-targeting siRNA pool (ON-TARGETplus siRNA SMARTpool; L-005077-00-0005, Dharmacon) or control siRNA (AllStars Negative Control, Qiagen) were transfected at 30 nM final concentration.

### High-content imaging analysis-guided SARS-CoV-2 infectivity assay

High-content imaging-guided SARS-CoV-2 infectivity assay was performed as previously described, with minor modifications [26]. Briefly, A549/ACE2 cells in suspension were infected with SARS-CoV-2 viruses at an MOI (multiplicity of infection) of 0.07 in serum-free DMEM/F12 medium with Glutamax^TM^ supplemented with 1X NEAA, at 37°C with 5% CO2 in a humidified incubator under gentle shaking. One hour post-infection, virus-containing medium was removed and replaced with complete medium, and when applicable, medium containing indicated compounds or DMSO, followed by overnight incubation at 37°C with 5% CO2 in a humidified incubator. Cells were then washed with PBS twice, fixed in 4% paraformaldehyde for 20 minutes at room temperature, and washed again with PBS twice and stored at 4 °C till immunofluorescence staining.

To detect virus-infected cells, cells were stained with anti-SARS-CoV-2 nucleocapsid (NC) monoclonal antibody (B46F) (1: 120 in 4% fetal bovine serum/PBS) at 4 °C overnight, followed by Alexa488-conjugated anti-mouse secondary antibody (1: 500 in PBS) mixed with 1 µg/ml DAPI at room temperature for 1-2h. Cells were washed by PBS thrice between each incubation. Finally, images of cells were acquired on an ImageXpress (Molecular Devices) microscope, and SARS-CoV-2 NC and/or DAPI-positive cells were identified and quantified using Cell Profiler software. For each assay condition per experiment, ≥ 3000 cells were analysed. Percentage of infection is calculated as the ratio between number of DAPI/SARS-CoV-2 NC double-positive cells and total number of DAPI-positive cells, which is further normalized to untreated, virus-infected samples.

When applicable, infectivity percentages were used to generate compound antiviral EC_50_ values *via* a nonlinear curve fitting model with variable slope (GraphPad Prism).

#### Cell cycle analysis

Cells in clear-bottomed 96-well plates (BD Falcon) were subsequently fixed with 4% paraformaldehyde in PBS for 20 minutes, permeabilized with 0.1% Triton X-100 in PBS for 30 minutes, blocked with 4% fetal bovine serum for 30 minutes, and finally stained with 1 µg/ml DAPI in PBS for 1 hour. Alternatively, cells were treated with 10 µM EdU for 30min before cells were washed twice with PBS followed by fixation, permeabilization, and blocking as described above. Azide-alkyne click reaction was then performed to conjugate EdU with Alexa Fluor 647 Azide (Invitrogen) fluorophore using the following reaction mix – 4mM CuSO4, 6nM Alexa Fluor 647 Azide, 10mM ascorbic acid in PBS. Cells were subsequently stained for DAPI in PBS as described above. All the incubations were performed at room temperature, and cells were washed with PBS twice between each incubation. Images of cells were acquired on an ImageXpress (Molecular Devices) microscope, and then analysed (nuclei counting, DAPI/EdU intensity measurements) with CellProfiler (Broad Institute) and data handled in Excel (Microsoft) and plotted in Prism 8 (GraphPad).

#### Intracellular nucleotide pool measurement

Intracellular nucleotide pool measurement was performed as described previously, with minor modifications [41, 42]. Briefly, A549/ACE2 cells at 70-80% confluency were treated with compounds of indicated concentrations or 1.2% DMSO for 6 hours at 37°C with 5% CO2 in a humidified incubator. Cells were then washed with TBS over ice, and then harvested in 700 μl ice-cold TCA solution (15% trichloroacetic acid, 30mM MgCl_2_) by using a cell scraper, before being snap frozen in liquid nitrogen and then stored at -80°C till nucleotide extraction and analysis via strong anion-exchange HPLC coupled with UV detection, which were performed as described previously [42].

## QUANTIFICATION AND STATISTICAL ANALYSIS

Statistical analysis was conducted using GraphPad Prism 9 software. Specific statistical test details, including test method, n values (number of independent experiments), number of technical repeats per independent experiment, and dispersion and precision measures, are indicated in the corresponding figure legends. Statistical significance is defined as p < 0.05, unless otherwise stated. Asterisk in figures signifies statistical significance (* for p ≤ 0.05, ** for p ≤ 0.01, *** for p ≤ 0.001, **** for p ≤ 0.0001).

## SUPPLEMENTARY ITEMS

**Figure S1 Supplemental information of UCK inhibition antagonized NHC-induced cell growth inhibition, related to Figure 1. A.** UCK inhibitor CPU dose-dependently antagonized the cytotoxic efficacy of NHC in multiple cell lines. Cells were treated with a dose-response matrix of NHC and CPU for 4 days before cell viability was determined using the resazurin reduction assay. Mean cell viability relative to DMSO control group ± SEM of n=2-3 independent experiments are shown, which were further curve fitted using a nonlinear curve fitting model with variable slope (GraphPad Prism) to generate the cell viability curves. **B-E**. Ponceau red staining images of Western blot membranes shown in Figure 1E, to demonstrate cell lysates containing equal amounts of total proteins were analysed. In B, same membrane later probed for UCK1 and UCK2, generating Western blot image shown in Figure 1E left panel; in C and D, membranes later probed for UCK1 (C) and UCK2 (D), generating Western blot image shown in Figure 1E right panel. Sample identities are shown in E.

**Figure S2 Supplemental information of DSF experiments on recombinant UCK proteins, incubated with nucleotides, related to Figure 2. A-B.** The non-substrate thymidine did not effectively engage recombinant UCK1, when applied up to 4.8mM in the DSF assay. Recombinant UCK1 (5 µM) was incubated with indicated concentrations of thymidine or DMSO, in the presence or absence of ATP, before protein thermal stabilities were determined using the DSF assay. In A, *left panel* displays melting curves of a representative experiment performed in quadruplicate, where mean fluorescence signals (solid line) ± SEM (dashed line) are shown; *right panel* displays negative derivative (-dRFU/dT) of the melting curves in the left panel, where mean negative derivative values (solid lines) ± SEM (dashed lines) are shown. Protein melting temperatures (Tm) were determined as the local minima of the negative derivative curves, and changes of Tm compared to ATP-only group are shown as ΔTm in B, where mean ΔTm ± SEM of n=4 independent experiments performed in triplicate to quadruplicate are shown. **C-F**. Melting profiles of recombinant UCK1 (C-D) and UCK2 (E-F) when incubated with NHC or C. Recombinant UCK proteins were incubated with increasing concentrations of NHC, C or DMSO, in the presence or absence of ATP, before protein thermal stabilities were xdetermined using the DSF assay. *Left panels*, melting curves of a representative experiment performed in quadruplicate, where mean fluorescence signals (solid line) ± SEM (dashed line) are shown; *right panel*, negative derivative (-dRFU/dT) of the melting curves in the left panel, where mean negative derivative values (solid lines) ± SEM (dashed lines) are shown. The curves are supplemental to Figure 2 E-F.

**Figure S3 Supplemental information of kinetic studies using the ADP Glo-coupled kinase activity assay, related to Figure 3. A-B.** Titration of UCK1 (A) and UCK2 (B) when using cytidine or NHC as the substrate. Recombinant UCK1 and UCK2 of varying concentrations were incubated with 200 µM or 100 µM substrates, respectively. Reactions were allowed to proceed for indicated periods before the addition of ADP Glo reagent and kinase detection reagent, with incubation of 1 hour between each addition. Luminescence measurement was then done on a Hidex plate reader. Mean luminescence signals ± SEM of independent experiments performed in triplicate are shown. **C-D**. Kinetic studies of phosphorylation of cytidine by recombinant UCK1 (C) and UCK2 (D), supplementary to Figure 3 A and B, respectively. The reaction was linear under the specified conditions for the duration of the experiment. Mean luminescence ± SEM of a representative experiment performed in triplicate are shown, which were subsequently used to determine V0 and reaction kinetic parameters. **E-F**. Kinetic studies of phosphorylation of NHC by recombinant UCK1 (E) and UCK2 (F), supplementary to Figure 3 C and D, respectively. The reaction was linear under the specified conditions for the duration of the experiment. Mean luminescence ± SEM of a representative experiment performed in triplicate are shown, which were subsequently used to determine V0 and reaction kinetic parameters.

**Figure S4 Supplemental information of UCK2 downregulation-induced loss of NHC antiviral efficacy, related to Figure 4. A-B.** The high-content imaging-based SARS-CoV-2 infectivity assay produced antiviral efficacies of NHC (A) and Remdesivir (B) that are comparable to past studies. A549/ACE2 cells were infected overnight with SARS-CoV-2 at an MOI of 0.07 in the presence of antivirals or the diluent control DMSO, before cells were fixed and stained for viral nucleocapsid protein (NC) and nuclei using NC-specific antibody and DAPI, respectively. Infectivity was estimated as the ratio of NC/DAPI double-positive cells over DAPI single-positive cells, and further normalized to DMSO-only control group. The resulting relative infectivity (%) data were then curve fitted using a nonlinear curve fitting model with variable slope (GraphPad Prism) to generate the antiviral EC_50_. The estimated infectivity agreed with mean NC intensity in cells stained positive for DAPI, showcasing the robustness of the automated cell profiler pipeline for identifying NC/DAPI double-positive cells. Mean infectivity relative to DMSO-only control group ± SEM, as well as mean NC intensity/cells/well, of a representative experiment performed in duplicate are shown. **C.** UCK2 knockdown did not affect A549/ACE2 cell cycle progression. A549/ACE2 cells were transfected with UCK2-specific (siUCK2) or non-targeting control (siNT) siRNA. Four days post-transfection, cells were fixed and stained for DAPI, followed by high-content imaging acquisition and DAPI signal analysis using CellProfiler. Frequency distribution analysis of DAPI signals were then done using GraphpadPrism. **D.** UCK2 knockdown did not affect A549/ACE2 cell proliferation. A549/ACE2 cells were transfected with UCK2-specific (siUCK2) or non-targeting control (siNT) siRNA. One day post-transfection, cells were re-seeded and allowed to proliferate for four days, before cell growth was determined via the confluence analysis function on a Tecan Spark® Cyto imaging cytometer. *Left panel*, mean cell confluence ± SEM of n=2 independent experiments; right panel, representative cell growth pictures used for confluence analysis. **E.** During the duration of viral infectivity assay, neither NHC (left panel) nor remdesivir (Rem, right panel), significantly affected the viability of siUCK2 or siNT-transfected A549/ACE2 cells. A549/ACE2 cells were transfected with siUCK2 or siNT siRNA. Three days post-transfection, cells were treated overnight with NHC (left panel), remdesivir (right panel), or the diluent control DMSO, at the same concentrations used for viral infectivity assay as shown in Figure 4 E-F. Cell viability was subsequently determined using resazurin reduction viability assay and normalized to DMSO-only control group. Mean relative viability ± SEM of n=2 independent experiments are shown. **F-G.** UCK2 knockdown significantly hampered intracellular NHC-TP pool, with no effect on NTP levels. A549/ACE2 cells were transfected with siUCK2 or siNT siRNA. One day post-transfection, cells were treated with 100 µM NHC before cell lysates were harvested, and intracellular nucleotide pools were measured via HPLC. In F, mean NTP levels per 10^6^ cells ± SEM of n=2 independent experiments, together with individual experiment values, are shown. In G, mean NHC-TP levels per 10^6^ cells ± SEM of n=2 independent experiments, together with individual experiment values, are shown. NHC-TP levels were further normalized to total NTP levels and the resulting NHC-TP/total NTP values ± SEM of n=2 independent experiments, together with individual experiment values, are also shown in G. Unpaired t-tests were performed across treatment groups – for NHC-TP levels per 10^6^ cells [siNT vs. siUCK2, p = 0.0137, t=8, df=2]; for NHC-TP/total NTP [siNT vs. siUCK2, p = 0.004, t=15, df=2], where asterisk signifies statistical significance (∗p ≤ 0.05, ∗∗p ≤ 0.01).

**Figure S5 Supplemental information of CPU-induced loss of NHC antiviral efficacy, related to Figure 5. A.** UCK inhibition via CPU did not affect A549/ACE2 cell proliferation, shown via confluence analysis. A549/ACE2 cells were seeded overnight and then treated with indicated concentrations of CPU or the diluent control DMSO for four days before cell growth was assessed using the confluence analysis function on a Tecan Spark® Cyto imaging cytometer. *Left panel*, mean cell confluence relative to DMSO-only group ± SEM of n=2 independent experiments; right panel, representative cell growth pictures used for confluence analysis. Dunnett’s multiple comparisons tests were performed across treatment groups [0 µM CPU vs. 22.1 µM CPU, p = 0.9762, q = 0.3173, DF = 4; 0 µM CPU vs. 75 µM CPU, p = 0.7345, q = 0.8690, DF = 4; 0 µM CPU vs. 256 µM CPU, p = 0.7345, q = 0.8690, DF = 4], where ns signifies statistical insignificance (p > 0.05). **B-C**. During the course of the 1-day viral infectivity assay, NHC and/or CPU did not affect cell viability (B) or cell cycle progression (C). A549/ACE2 cells were treated overnight with a concentration matrix of NHC and CPU, at the same concentrations as in viral infectivity assay shown in Figure 5. Following the treatment, in B, cell viabilities were determined using the resazurin reduction viability assay. Relative viability normalized to cells treated with DMSO only ± SEM of n=2 independent experiments are shown. In C, cell cycle was determined following the treatment with highest concentrations of NHC and/or CPU, where DAPI-stained cells were imaged by high-content imaging followed by DAPI intensity analysis via CellProfiler. *Left panel*, DAPI-intensity histograms with gating strategy of a representative experiment; *right panel*, mean percentages of cells in different cell cycle phases ± SEM of n=2 independent experiments, together with individual experiment values, are shown. **D-E**. CPU significantly hampered intracellular NHC-TP pool, with no effect on NTP levels. A549/ACE2 cells were pre-treated with 256 µM CPU or DMSO for an hour before being further treated with 100 µM NHC or DMSO for 6 hours. Cell lysates were then harvested, and intracellular nucleotide pools were measured via HPLC. In D, mean NTP levels per 10^6^ cells ± SEM of n=3 independent experiments, together with individual experiment values, are shown. In E, mean NHC-TP levels per 10^6^ cells ± SEM of n=3 independent experiments, together with individual experiment values, are shown. NHC-TP levels were further normalized to total NTP levels and the resulting NHC-TP/total NTP values ± SEM of n=3 independent experiments, together with individual experiment values, are also shown in E. Ratio paired t-tests were performed across treatment groups – for NHC-TP levels per 10^6^ cells [0 µM CPU vs. 256 µM CPU, p = 0.0025, t=20.09, df=2]; for NHC-TP/total NTP [0 µM CPU vs. 256 µM CPU, p = 0.0066, t=12.23, df=2], where asterisk signifies statistical significance (∗∗p ≤ 0.01). **F.** During the course of the 1-day viral infectivity assay, remdesivir (Rem) and/or CPU did not affect cell viability. A549/ACE2 cells were treated overnight with Rem and/or CPU, at the same concentrations as in viral infectivity assay shown in Figure 5, which was then followed by resazurin reduction viability assay to determine cell viability. Relative viability normalized to cells treated with DMSO only ± SEM of n=2 independent experiments are shown.

**Figure S6 Supplemental information of CPU-induced reduction of the selectivity index of NHC, related to Figure 6. A-B.** Four day-treatment with NHC induced S phase collapse and G1 arrest in A549/ACE2 cells. A549/ACE2 cells were treated with increasing concentrations of NHC for four days before cell cycles were analyzed via immunofluorescence staining for EdU incorporation and DAPI followed by high-content imaging. In A, representative plots with gating strategy illustrated. In B, percentage of cells in each cell cycle (*left panel*) and mean integrated EdU signal of cells in S phase (*right panel*) of a representative experiment, illustrating that NHC dose-dependently induced collapse of EdU-incorporating S phase cells concurrent to G1 arrest. **C.** Viral infectivity inhibition curves of SARS-CoV-2 infection in A549/ACE2 cells, and the viability curves of A549/ACE2 cells generated using the same dose-response matrix of CPU and NHC. In the viral infectivity assay, A549/ACE2 cells were infected with SARS-CoV-2 in the absence or presence of CPU and/or NHC. One day post-infection, viral infectivity was determined via staining the cells with viral nucleocapsid protein followed by high-content imaging analysis. Mean relative infectivity normalized to the DMSO-only group ± SEM of n=4 independent experiments performed in duplicate are shown. Meanwhile the effect of NHC on cell viability was determined with the same NHC/CPU matrix but a prolonged four-day drug treatment, followed by resazurin viability reduction assay. Mean viability relative to DMSO group ± SEM of n=2-4 independent experiments are shown. Both viability and infectivity data were further curve fitted using a nonlinear curve fitting model with variable slope (GraphPad Prism) to generate the antiviral and growth inhibitory EC_50_ values of NHC, which were further used to calculate selectivity index as shown in Figure 6D.

**Figure S7 Supplemental information of recombinant UCK1 protein production and purification.** Purified recombinant UCK1 protein (4 µg) was subject to SDS-PAGE followed by Coomassie brilliant blue staining, demonstrating protein purity.

## REFERENCES

1. Van Rompay, A.R., et al., Phosphorylation of uridine and cytidine nucleoside analogs by two human uridine-cytidine kinases. Mol Pharmacol, 2001. 59(5): p. 1181–6.

2. Fu, Y., et al., The Metabolic and Non-Metabolic Roles of UCK2 in Tumor Progression. Frontiers in Oncology, 2022. **Volume 12 -** 2022.

3. Drake, J.C., R.G. Stoller, and B.A. Chabner, Characteristics of the enzyme uridine-cytidine kinase isolated from a cultured human cell line. Biochemical Pharmacology, 1977. 26(1): p. 64–66.

4. Van Rompay, A.R., M. Johansson, and A. Karlsson, Phosphorylation of Deoxycytidine Analog Monophosphates by UMP-CMP Kinase: Molecular Characterization of the Human Enzyme. Molecular Pharmacology, 1999. 56(3): p. 562–569.

5. Sarkisjan, D., et al., The Cytidine Analog Fluorocyclopentenylcytosine (RX-3117) Is Activated by Uridine-Cytidine Kinase 2. PLoS One, 2016. 11(9): p. e0162901.

6. Valencia, A., et al., Expression of nucleoside-metabolizing enzymes in myelodysplastic syndromes and modulation of response to azacitidine. Leukemia, 2014. 28(3): p. 621–628.

7. Gu, X., et al., Decitabine- and 5-azacytidine resistance emerges from adaptive responses of the pyrimidine metabolism network. Leukemia, 2021. 35(4): p. 1023–1036.

8. Sripayap, P., et al., Mechanisms of resistance to azacitidine in human leukemia cell lines. Experimental Hematology, 2014. 42(4): p. 294–306.e2.

9. Sripayap, P., et al., Mechanisms of resistance to azacitidine in human leukemia cell lines. Exp Hematol, 2014. 42(4): p. 294–306 e2.

10. Wahl, A., et al., SARS-CoV-2 infection is effectively treated and prevented by EIDD-2801. Nature, 2021. 591(7850): p. 451-457.

11. Ehteshami, M., et al., Characterization of β-d-N(4)-Hydroxycytidine as a Novel Inhibitor of Chikungunya Virus. Antimicrob Agents Chemother, 2017. 61(4).

12. Stuyver, L.J., et al., Ribonucleoside analogue that blocks replication of bovine viral diarrhea and hepatitis C viruses in culture. Antimicrob Agents Chemother, 2003. 47(1): p. 244–54.

13. Yoon, J.J., et al., Orally Efficacious Broad-Spectrum Ribonucleoside Analog Inhibitor of Influenza and Respiratory Syncytial Viruses. Antimicrob Agents Chemother, 2018. 62(8).

14. Reynard, O., et al., Identification of a New Ribonucleoside Inhibitor of Ebola Virus Replication. Viruses, 2015. 7(12): p. 6233–40.

15. Agostini, M.L., et al., Small-Molecule Antiviral β-d-N(4)-Hydroxycytidine Inhibits a Proofreading-Intact Coronavirus with a High Genetic Barrier to Resistance. J Virol, 2019. 93(24).

16. Kabinger, F., et al., Mechanism of molnupiravir-induced SARS-CoV-2 mutagenesis. Nature Structural & Molecular Biology, 2021. 28(9): p. 740–746.

17. Zhou, S., et al., β-d-N4-hydroxycytidine Inhibits SARS-CoV-2 Through Lethal Mutagenesis But Is Also Mutagenic To Mammalian Cells. The Journal of Infectious Diseases, 2021. 224(3): p. 415–419.

18. Xu, Z., et al., Uridine–cytidine kinase 2 potentiates the mutagenic influence of the antiviral β-d-N4-hydroxycytidine. Nucleic Acids Research, 2023.

19. Feng, J.Y., Addressing the selectivity and toxicity of antiviral nucleosides. Antivir Chem Chemother, 2018. 26: p. 2040206618758524.

20. Lim, M.I., et al., Cyclopentenyluridine and cyclopentenylcytidine analogs as inhibitors of uridine-cytidine kinase. Journal of Medicinal Chemistry, 1984. 27(12): p. 1536–1538.

21. Karle, J.M. and R.L. Cysyk, Regulation of pyrimidine biosynthesis in cultured L1210 cells by 3-deazauridine. Biochemical Pharmacology, 1984. 33(23): p. 3739–3742.

22. Moriconi, W.J., M. Slavik, and S. Taylor, 3-Deazauridine (NSC 126849): an interesting modulator of biochemical response. Invest New Drugs, 1986. 4(1): p. 67–84.

23. Kang, G.J., et al., Cyclopentenylcytosine triphosphate. Formation and inhibition of CTP synthetase. J Biol Chem, 1989. 264(2): p. 713–8.

24. Toots, M., et al., Quantitative efficacy paradigms of the influenza clinical drug candidate EIDD-2801 in the ferret model. Translational Research, 2020. 218: p. 16–28.

25. Okesli-Armlovich, A., et al., Discovery of small molecule inhibitors of human uridine-cytidine kinase 2 by high-throughput screening. Bioorganic & Medicinal Chemistry Letters, 2019. 29(18): p. 2559–2564.

26. Zhang, S.M., et al., NUDT15-mediated hydrolysis limits the efficacy of anti-HCMV drug ganciclovir. Cell Chemical Biology, 2021.

27. Chou, S.W. and K.M. Scott, Rapid quantitation of cytomegalovirus and assay of neutralizing antibody by using monoclonal antibody to the major immediate-early viral protein. Journal of clinical microbiology, 1988. 26(3): p. 504–507.

28. Akinci, E., et al., Elucidation of remdesivir cytotoxicity pathways through genome-wide CRISPR-Cas9 screening and transcriptomics. bioRxiv, 2020.

29. Lim, S.-Y., et al., Anti-SARS-CoV-2 Activity of Adamantanes In Vitro and in Animal Models of Infection. COVID, 2022. 2(11): p. 1551–1563.

30. Cox, R.M., J.D. Wolf, and R.K. Plemper, Therapeutically administered ribonucleoside analogue MK-4482/EIDD-2801 blocks SARS-CoV-2 transmission in ferrets. Nat Microbiol, 2021. 6(1): p. 11–18.

31. Cox, R.M., et al., Oral prodrug of remdesivir parent GS-441524 is efficacious against SARS-CoV-2 in ferrets. Nature Communications, 2021. 12(1): p. 6415.

32. Grant, S., K. Bhalla, and M. Gleyzer, Effect of Uridine on Response of 5-Azacytidine-resistant Human Leukemic Cells to Inhibitors of de Novo Pyrimidine Synthesis1. Cancer Research, 1984. 44(12_Part_1): p. 5505-5510.

33. Liu, J., et al., SARS-CoV-2 cell tropism and multiorgan infection. Cell Discovery, 2021. 7(1): p. 17.

34. Maas, B.M., et al., Molnupiravir: Mechanism of action, clinical, and translational science. Clin Transl Sci, 2024. 17(2): p. e13732.

35. Iwamoto, M., et al., Assessment of pharmacokinetics, safety, and tolerability following twice-daily administration of molnupiravir for 10 days in healthy participants. Clinical and Translational Science, 2023. 16(10): p. 1947–1956.

36. Girish, V., et al., Oncogene-like addiction to aneuploidy in human cancers. Science, 2023. 381(6660): p. eadg4521.

37. Bonagas, N., et al., Pharmacological targeting of MTHFD2 suppresses acute myeloid leukemia by inducing thymidine depletion and replication stress. Nat Cancer, 2022. 3(2): p. 156–172.

38. Zhang, S.M., et al., Development of a chemical probe against NUDT15. Nature Chemical Biology, 2020.

39. Zhang, S.M., et al., Identification and evaluation of small-molecule inhibitors against the dNTPase SAMHD1 via a comprehensive screening funnel. iScience, 2024: p. 108907.

40. Yadav, B., et al., Searching for Drug Synergy in Complex Dose-Response Landscapes Using an Interaction Potency Model. Comput Struct Biotechnol J, 2015. 13: p. 504–13.

41. Rentoft, M., et al., Heterozygous colon cancer-associated mutations of SAMHD1 have functional significance. Proc Natl Acad Sci U S A, 2016. 113(17): p. 4723–8.

42. Ranjbarian, F., et al., Isocratic HPLC analysis for the simultaneous determination of dNTPs, rNTPs and ADP in biological samples. Nucleic Acids Res, 2022. 50(3): p. e18.

