## Supplementary information for "Uridine cytidine kinases dictate the therapeutic response of molnupiravir via its bioactivation"

Figure S1

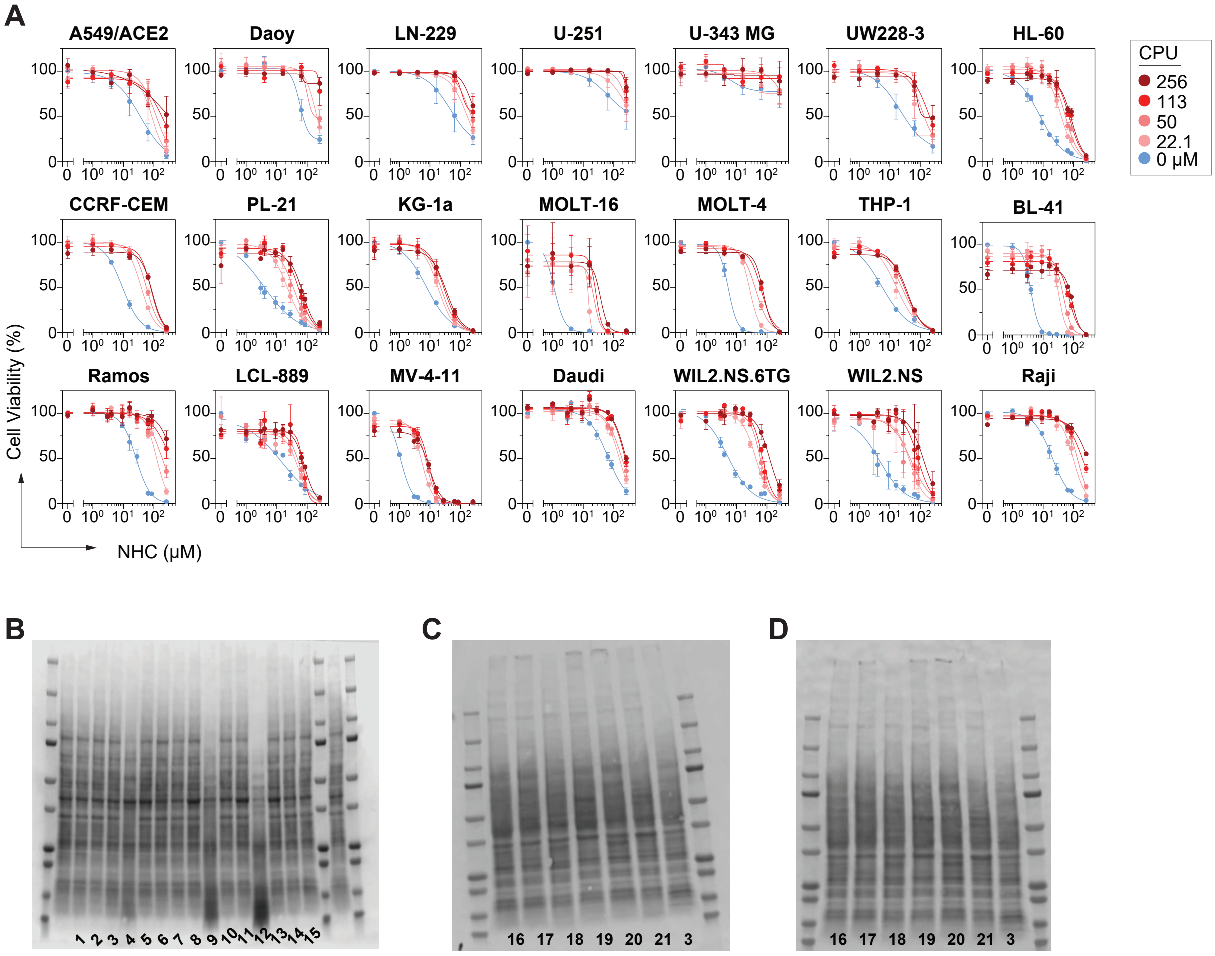


E

| | | | | | | | | | |
|---|---|---|---|---|---|---|---|---|---|
| 1 | MOLT-4 | 6 | KG-1a | 11 | WIL2.NS.6TG | 16 | UW228-3 | 21 | A549/ACE2 |
| 2 | CCRF-CEM | 7 | Ramos | 12 | PL-21 | 17 | Daoy | | |
| 3 | HL-60 | 8 | LCL-889 | 13 | WIL2.NS | 18 | U-343 MG | | |
| 4 | THP-1 | 9 | MV-4-11 | 14 | Raji | 19 | U-251 MG | | |
| 5 | MOLT-16 | 10 | Daudi | 15 | BL-41 | 20 | LN-229 | | |

Figure S2

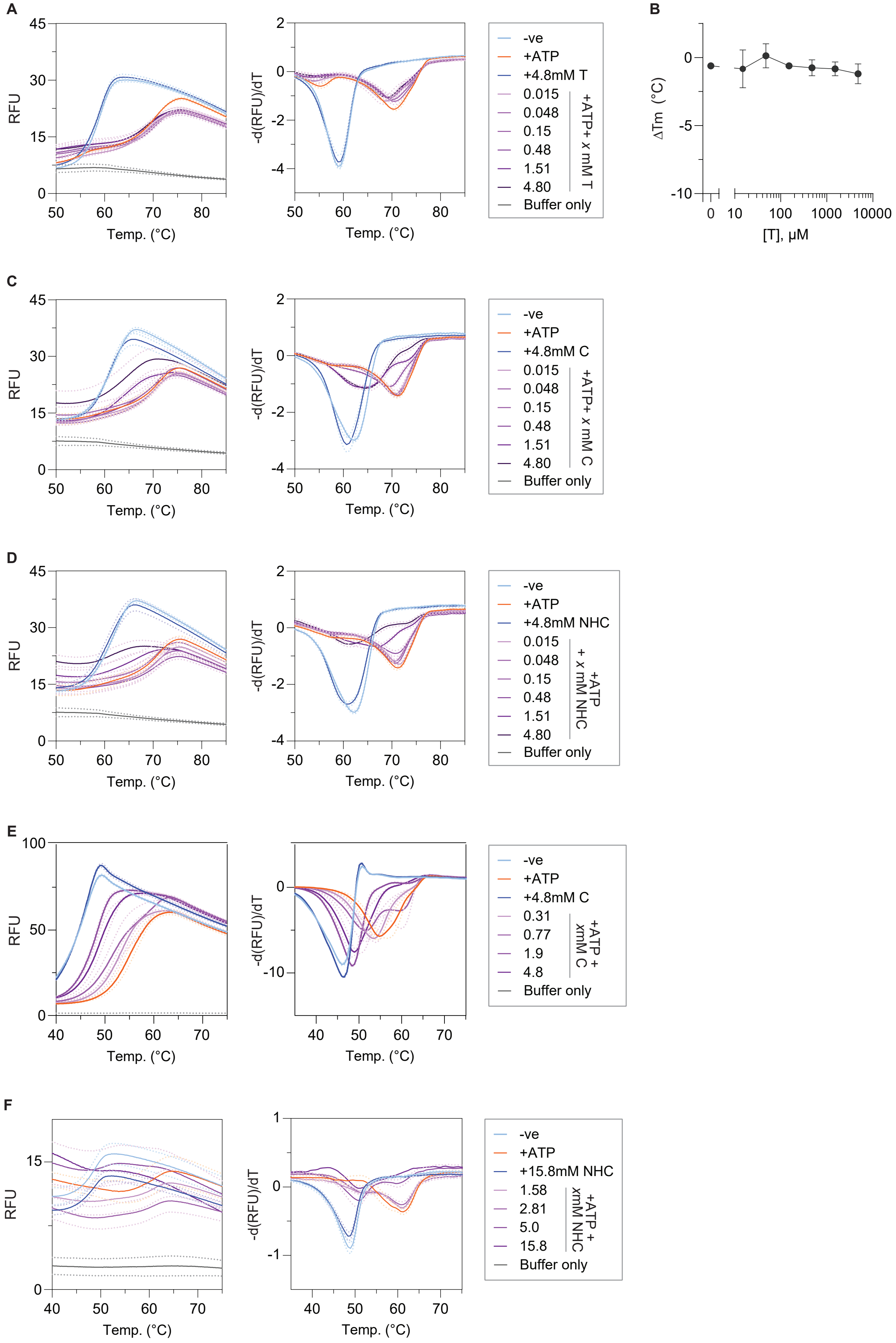
A
RFU
Temp. (°C)
-d(RFU)/dT
-ve
+ATP
+4.8mM T
0.015
0.048
0.15
0.48
1.51
4.80
+ATP+ x mM T
Buffer only
B
ΔTm (°C)
[T], μM
C
-ve
+ATP
+4.8mM C
0.015
0.048
0.15
0.48
1.51
4.80
+ATP+ x mM C
Buffer only
D
-ve
+ATP
+4.8mM NHC
0.015
0.048
0.15
0.48
1.51
4.80
+ATP + x mM NHC
Buffer only
E
-ve
+ATP
+4.8mM C
0.31
0.77
1.9
4.8
+ATP + xmM C
Buffer only
F
-ve
+ATP
+15.8mM NHC
1.58
2.81
5.0
15.8
+ATP + xmM NHC
Buffer only

Figure S3

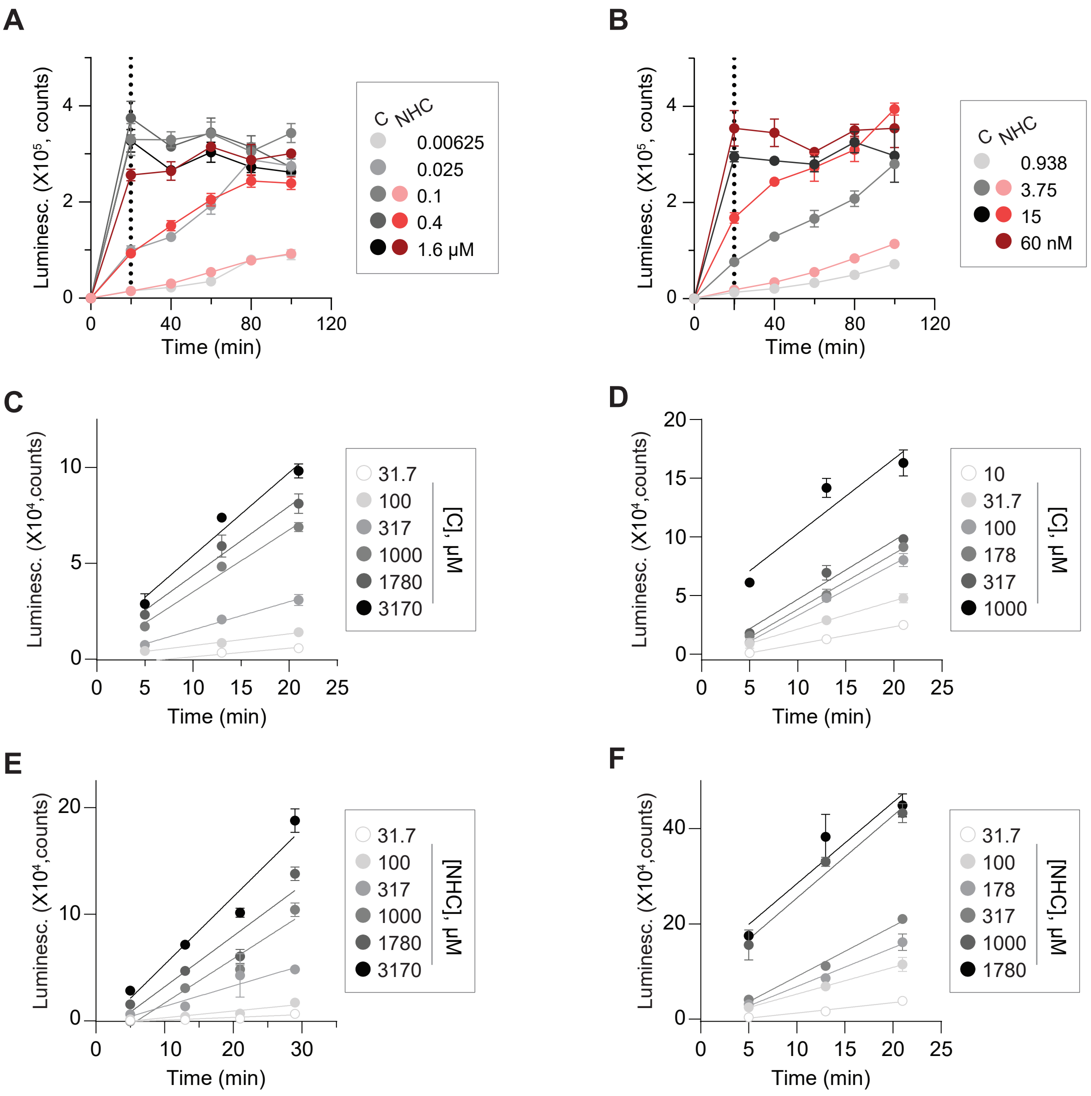
A
B
C
D
E
F
Luminesc. (X10⁵, counts)
Time (min)
C NHC
0.00625
0.025
0.1
0.4
1.6 µM
0.938
3.75
15
60 nM
Luminesc. (X10⁴,counts)
31.7
100
317
1000
1780
3170
[C], µM
10
178
[NHC], µM

### Figure S4

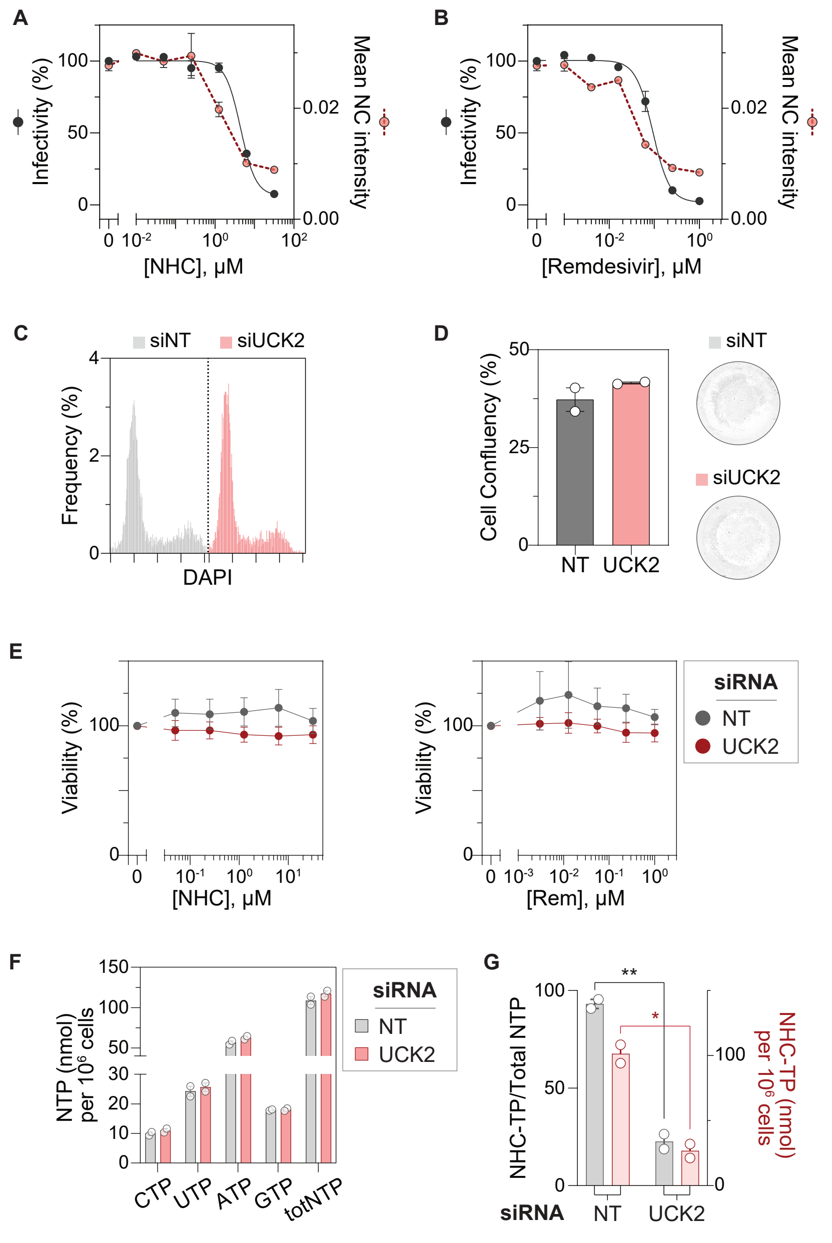
A
Infectivity (%)
Mean NC intensity
[NHC], µM
B
[Remdesivir], µM
C
siNT
siUCK2
Frequency (%)
DAPI
D
Cell Confluency (%)
NT
UCK2
E
Viability (%)
siRNA
[NHC], µM
[Rem], µM
F
NTP (nmol) per 10^6 cells
CTP
UTP
ATP
GTP
totNTP
G
NHC-TP/Total NTP
NHC-TP (nmol) per 10^6 cells
**
*

Figure S5

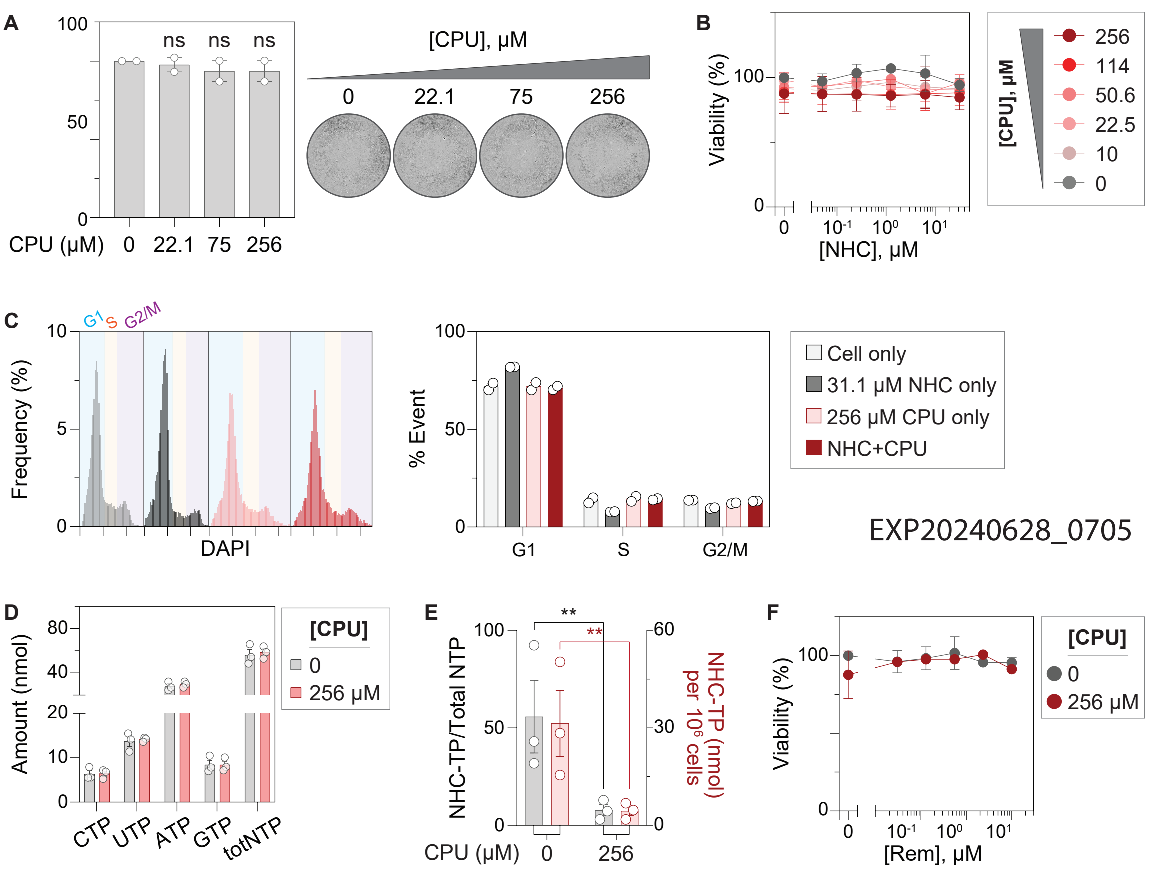
A
ns
ns
ns
CPU (μM) 0 22.1 75 256
[CPU], μM
0 22.1 75 256
B
Viability (%)
[NHC], μM
[CPU], μM
256
114
50.6
22.5
10
0
C
G1 S G2/M
Frequency (%)
DAPI
% Event
G1 S G2/M
Cell only
31.1 μM NHC only
256 μM CPU only
NHC+CPU
EXP20240628_0705
D
Amount (nmol)
CTP UTP ATP GTP totNTP
[CPU]
0
256 μM
E
NHC-TP/Total NTP
NHC-TP (nmol) per 10^6 cells
CPU (μM) 0 256
F
Viability (%)
[Rem], μM
[CPU]
0
256 μM

Figure S6

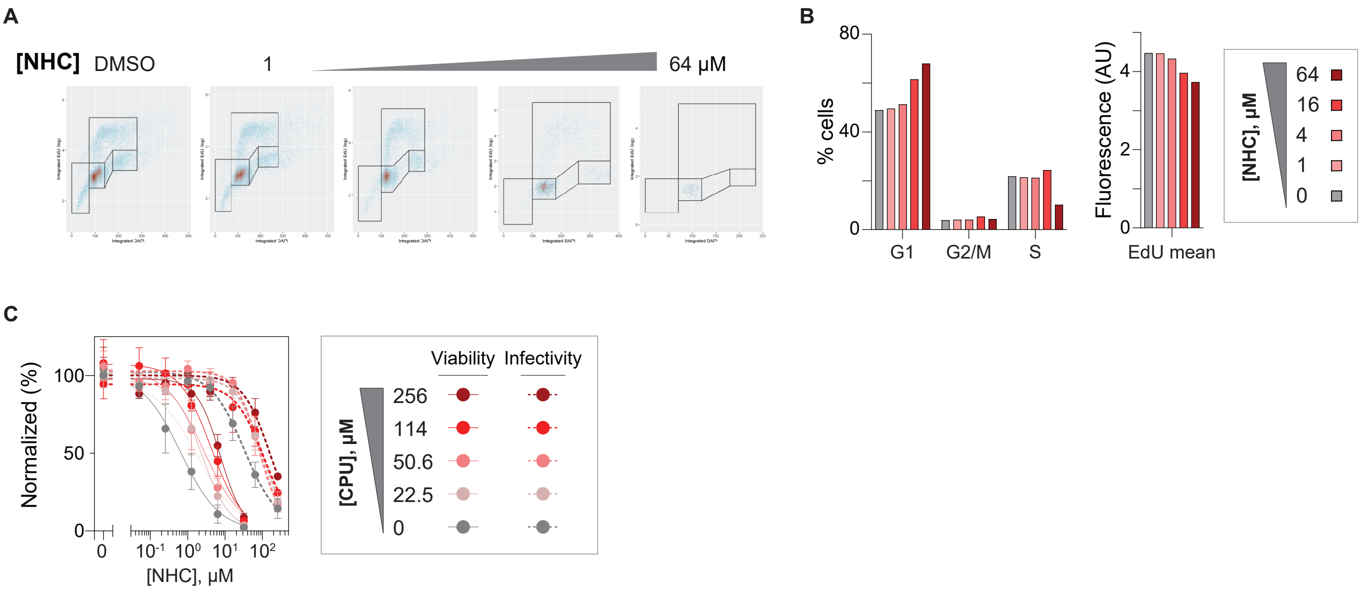
A
[NHC] DMSO
1
64 μM
B
80
40
0
% cells
G1
G2/M
S
Fluorescence (AU)
4
2
0
EdU mean
[NHC], μM
64
16
4
1
0
C
Normalized (%)
100
50
0
0
10-1 10⁰ 10¹ 10²
[NHC], μM
[CPU], μM
Viability
Infectivity
256
114
50.6
22.5
0

Figure S7

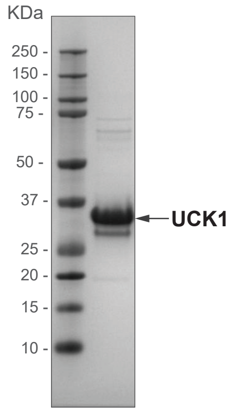

Uncropped WB blots

Fig1E

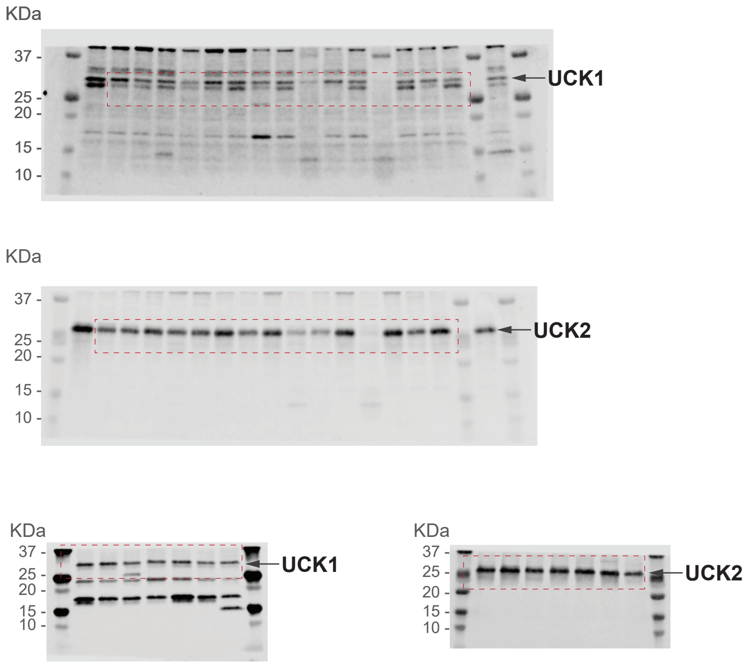


Fig4A

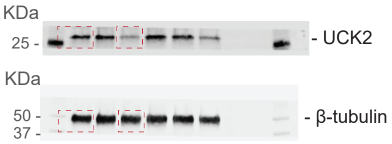
